# Hydrodynamic and anthropogenic disturbances co-shape microbiota rhythmicity and community assembly within intertidal groundwater-surface water continuum

**DOI:** 10.1101/2022.11.06.515374

**Authors:** Ze Zhao, Lu Zhang, Guoqing Zhang, Han Gao, Xiaogang Chen, Ling Li, Feng Ju

## Abstract

Tidal hydrodynamics drive the groundwater-seawater exchange and shifts in microbiota structure in the coastal zone. However, how the coastal water microbiota structure and assembly patterns respond to periodic tidal fluctuations and anthropogenic disturbance remain unexplored in the intertidal groundwater-surface water (GW-SW) continuum, although it affects biogeochemical cycles and coastal water quality therein. Here, through hourly time-series sampling in the saltmarsh tidal creek, rhythmic patterns of microbiota structure in response to daily and monthly tidal fluctuations in intertidal surface water are disentangled for the first time. The similarity in archaeal community structures between groundwater and ebb-tide surface water (R^2^=0.06, *p*=0.2) demonstrated archaeal transport through groundwater discharge, whereas multi-source transport mechanisms led to unique bacterial biota in ebb-tide water. Homogeneous selection (58.6%-69.3%) dominated microbiota assembly in the natural intertidal GW-SW continuum and the presence of 157 rhythmic ASVs identified at ebb tide and 141 at flood tide could be attributed to different environmental selection between groundwater and seawater. For intertidal groundwater in the tidal creek affected by anthropogenically contaminated riverine inputs, higher microbial diversity and shift in community structure were primarily controlled by increased co-contribution of dispersal limitation and drift (jointly 57.8%) and enhanced microbial interactions. Overall, this study fills the knowledge gaps in the tide-driven water microbial dynamics in coastal transition zone and the response of intertidal groundwater microbiota to anthropogenic pollution of overlying waters. It also highlights the potential of microbiome analysis in enhancing coastal water quality monitoring and identifying anthropogenic pollution sources (e.g., aquaculture pathogenic *Vibrio*) through the detection of rhythmic microbial variances associated with intertidal groundwater discharge and seawater intrusion.

## 1. Introduction

Coastal ecosystems (e.g., salt marshes and mangroves) play a critical role in global climate regulation [1] and biogeochemical cycles [2, 3], yet they are currently facing a multitude of local and global challenges caused by climate change and human activities (e.g., pollution). In addition to the well-known river-runoff input and atmosphere deposition, submarine groundwater discharge (SGD) or porewater exchange from coastal aquifers contributes to nutrients and microbial loadings onto the coastal ecosystems, thus affecting the microbially-driven biogeochemical cycling and water quality within the groundwater-seawater mixing system [4–6]. Most existing studies have focused on microbiota structure in only one compartment of the intertidal zone, such as coastal sediment [7], groundwater or coastal seawater [8–11]. However, microbiota dynamics in the intertidal groundwater-surface water (GW-SW) continuum, especially in the context of the high structural variability and complexity impacted by bidirectional water exchange, remain poorly explored. Moreover, although periodic disturbances (e.g., tidal fluctuation) are revealed to influence the microbiota structure in various environments [12, 13], high-resolution profiling of microbiota dynamics is lacking. Hourly time-series sampling, e.g., during daily or monthly tidal cycles, would help to resolve microbiota dynamic patterns and assembly rules in the intertidal aquatic continuum and further identify key processes and factors governing the microbiota structure.

Hydrodynamic disturbance can cause steep variations in physicochemical conditions over short spatial or temporal scales, and thus are regarded as a critical factor controlling microbiota assembly and biogeochemical processes in coastal sediments [12]. Various environmental factors (e.g., salinity, pH, temperature and nutrient [8, 14–17]) are implicated in exerting selective pressures on the intertidal groundwater or coastal seawater microbiome. The results are broadly consistent with the niche-based theory, which asserts that deterministic processes such as environmental filtering and biotic interactions largely control the patterns of microbiota composition and distribution [18]. In addition to deterministic mechanisms, stochastic processes, for instance, dispersal and ecological drift, also contribute to microbial assembly. According to the well-established community assembly theory [19, 20], microbiota structure and succession are controlled by four fundamental ecological processes: selection (homogeneous selection and heterogeneous selection), dispersal (homogenizing dispersal and dispersal limitation), drift and diversification, which vary in their degree of determinism and stochasticity [21] and the relative importance of the first three can be quantified by the null model [22, 23]. Until now, only a few studies have reported on the microbiota assembly patterns in the intertidal sediment [17, 24, 25] or coastal seawater [26], but the relative contribution of ecological processes greatly varied across studies. Comparatively, microbiota assembly processes have never been quantified in the intertidal GW-SW continuum, especially under co-effects of hydrodynamic and anthropogenic disturbances.

Deterministic processes, including environmental or habitat filtering and biotic interactions, are treated as joint selection forces in the community assembly and interact dynamically to drive the observed community patterns [27, 28]. However, their respective contribution cannot be parsed out based on the quantitative approach of the null model [23]. Instead, deterministic processes and ecological relationships can be interactively and topologically explored in a microbial co-occurrence or molecular ecological network, which is fundamental for predicting microbial species interactions and niche patterns in both natural [29, 30] and engineered [31–33] ecosystems. Previous studies have reported that enhanced seawater intrusion weakened the complexity of microbial co-occurrence associations in intertidal sediment [24], whereas it decreased the network size but caused higher compactness of groundwater microbial interactions [10]. However, microbial co-occurrence patterns in the intertidal GW-SW continuum and their dynamics under periodic tidal fluctuations and increasing anthropogenic impacts (e.g., land-sourced pollutants) are almost blank. Therefore, complementing null model with community-wide co-occurrence patterns will enable more complete insights into the ecological processes shaping microbial community diversity in the context of the determinism-versus-stochasticity dichotomy.

To explore the tide-driven dynamics of microbiota structure and community assembly mechanisms in an intertidal aquatic continuum, we collected intertidal groundwater and hourly time-series surface water in a natural saltmarsh tidal creek (NT) located in Jiangsu Province, China (Fig. 1A), where mudflats are estimated to cover more than 90% of the 954-km Jiangsu coast [34]. Meanwhile, a geographically adjacent but human-influenced tidal creek (HT) was chosen and compared with NT to resolve the response of intertidal groundwater microbiota to anthropogenic pollution of overlying waters. The objectives of this study are: (i) to investigate the temporal dynamics of microbiota structure in intertidal surface water with daily and monthly tidal fluctuations; (ii) to explore the processes and mechanisms involved in microbiota assembly in intertidal GW-SW continuum under tidal fluctuations; and (iii) to elucidate the effects of anthropogenically contaminated river input on the spatial distribution and community assembly of intertidal groundwater microbiota. This study will provide the first and in-depth understanding of microbiota dynamics and assembly mechanisms of the intertidal water co-shaped by hydrodynamic and anthropogenic disturbances in the coastal ecosystem.

**Figure 1.**
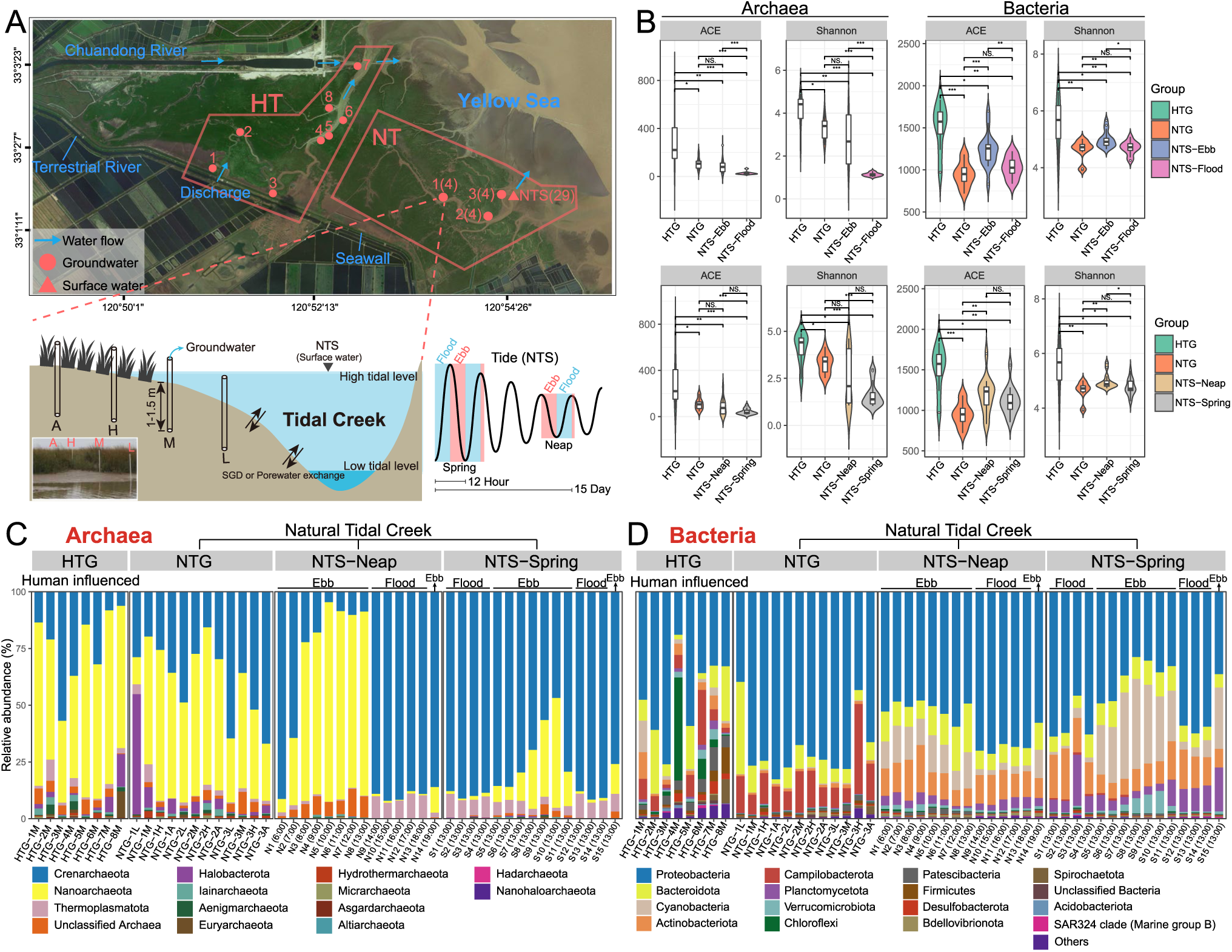
Sampling and overview of archaeal and bacterial communities in tidal creek groundwater and surface water. (A) Map of human-influenced (HT) and natural (NT) salt marsh tidal creeks sampled in this study. Red dots in the graph above show the sampling sites of intertidal groundwater and red triangle marks the times series station in NT for collecting surface water samples (NTS). Blue arrows show the water flow direction of tidal creek and Chuandong River, as well as the Terrestrial River water input upstream of HT. Groundwater samples from NT were collected at low (L), mid (M), high (H) and above high (A) tidal marks along three sampling transect of NT, whereas groundwater samples from HT were all collected at mid (M) tidal marks, as shown in the bottom-left graph. The blue and pink rectangles in the bottom-right graph mark the time periods for collecting time-series flood and ebb tide surface water, respectively, during neap tide and spring tide. (B) Alpha diversity shown as ACE and Shannon indices. The significance level in the median differences across groups is statistically checked using wilcoxon rank-sum tests. Significance is expressed as \*\*\**p* < 0.001, \*\**p* < 0.01, \**p* < 0.05, NS-non significant. (C) The composition of all phylum-level archaeal communities. (D) The composition of phylum-level bacterial communities. Top 16 most abundant phylum are shown and other phylum are classified as ‘Others’.

## 2. Materials and Methods

### 2.1. Field sample collection and physicochemical analysis

Samples were collected from the Saltmarsh Tidal Creeks located at Dafeng Milu National Nature Reserve, Yancheng City, Jiangsu Province, China, in August 2020 (Fig, 1A, Table S1). The reserve is located in Yancheng Coastal Wetland, which is the largest coastal wetlands nature reserve in China as listed in Ramsar Sites Information Service. Conducing microbial research here is representative and significant. Twelve groundwater samples were collected at low (L), mid (M), high (H) and above high (A) tidal marks along three sampling transects of the natural tidal creek (Natural tidal creek groundwater, NTG-1, NTG-2 and NTG-3). Eight groundwater samples were collected at mid (M) tidal marks of 8 sampling sites in the human-influenced tidal creek (Human-influenced tidal creek groundwater, HTG-1 to HTG-8). All groundwater samples were collected at the lowest surface water level during ebb tide, at a depth of 1.0-1.5 m. The outlet of HT is connected to the estuary of Chuandong River and the estuarine water flows into the creek at flood tide. Besides, the Terrestrial River upstream of HT discharges water into the creek at ebb tide, which is controlled by a floodgate. The upstream of Terrestrial River flows through human living (Yancheng City) and aquaculture area (bottom left of the map in Fig. 1A). Higher concentration of antibiotics for human and aquaculture use (0.66-3.07 ng/L of Metronidazole and 7.32-24.57 of Roxithromycin) and human-use-only Metformin (5.89-19.35) and sweetener (150.25-582.62), were detected in HT surface water compared with in NT surface water (0.20-1.11, 0-1.42, 0.49-2.12 and 64.06-128.21 ng/L) (Table S2), indicating serious water pollution in the HT (e.g., drug and nutrient) due to aquaculture wastewater and domestic sewage discharge. On the contrary, the downstream mouth of NT is far away from the Chuandong River estuary (about 4.5 km) and there is no terrestrial surface water input along the tidal creek, so it is considered a natural tidal creek under less anthropogenic influence.

To investigate the transport and dynamics of microbial communities impacted by bidirectional exchange of groundwater and coastal water during tidal fluctuations, surface water samples were collected hourly for 14-15 hours from the Times Series Station downstream of the NT (Natural tidal creek surface water, NTS) to monitor a complete phase of ebb and flood tide (Fig. 1A). This monitoring was conducted once at neap tide (N1-N14 on August 12^th^) and once at spring tide (S1-S15 on August 6^th^-7^th^) to investigate the difference in the intensity of tidal disturbances. Each water sample (300-1400 ml) was filtered on-site through 0.2 μm pore size polycarbonate filters (Millipore, USA) for about 30 minutes. The filters were transported to the laboratory with dry ice in 24 h and stored at −20 until DNA extraction. Physiochemical properties, including water depth, temperature (temp), pH, salinity, dissolved oxygen (DO), dissolved organic and inorganic carbon (DOC and DIC), and ^222^Rn (groundwater isotope tracer) of water samples were measured according to our earlier publication [35] (detailed in Supplementary Text S1 and Fig. S1).

In total, 49 water samples were grouped according to their sources and sampling times: (i) intertidal groundwater collected from human-influenced (HTG, 8) and natural tidal creek (NTG, 12); (ii) time-series surface water samples grouped according to two different grouping rules, namely NTS-Neap (14) and NTS-Spring (15) collected during neap and spring tide, or NTS-Ebb (17) and NTS-Flood (12) collected during ebb and flood tide designated by the *in-situ* observation of water depth.

### 2.2. DNA extraction, qPCR and 16S rRNA gene sequencing

Total genomic DNA was extracted using DNeasy PowerSoil Kit (Qiagen, Germany). The V4-V5 regions of prokaryotic 16S rRNA genes were amplified using primers 515F and 926R targeting both bacteria and archaea (recommended by the Earth Microbiome Project) [36], and sequenced on the Illumina NovaSeq 6000 platform with 250-bp paired-end reads at the Magigene Biotechnology Co. Ltd (Guangzhou, China). In addition, real-time quantitative PCR (qPCR) was performed in technical triplicates to quantify the 16S rRNA gene copies (as a proxy for prokaryotic biomass) in water samples using primers 515F and 926R on a Jena qTOWER^3^G (Germany) (detailed in Supplementary Text S2).

The raw sequencing data were processed following the QIIME2 pipeline [37] (detailed in Text S3). In brief, forward and reverse reads were trimmed to 240 and 220 base pairs and low-quality reads were removed using Trimmomatic [38]. All reads were then subjected to de-noising using the DADA2 pipeline with default parameters [39]. Taxonomic assignment of amplicon sequence variant (ASVs) was conducted using the Silva v138.1 database. All the following statistical analysis and visualization were performed with R software (4.0.5). The outputs from QIIME2, including table of amplicon sequence variants (ASVs), taxonomy and tree files were processed by R package phyloseq [40], MicrobiotaProcess and microeco [41]. Briefly, ACE and Shannon indices were used to evaluate the alpha-diversity of microbial communities. The ASV table was transformed to relative abundance using phyloseq function *transform_sample_counts.* Non-metric multidimensional scaling (NMDS) analysis and permutational multivariate analysis of variance (PERMANOVA) was performed based on weighted UniFrac distance. To test whether each of the ASVs in time-series surface water displayed rhythmic patterns in response to tidal fluctuations, that is, recurrent changes over time, we performed Lomb Scargle periodogram (LSP, as implemented in the lomb package) analysis for relative abundance of ASVs in NTS-Neap and NTS-Spring, respectively. The method has previously been used for testing the seasonality in marine microbial communities [42]. Briefly, we obtain density distribution of each period for each ASV, showing which is the most recurrent period for each ASV. Peak normalized power (PNmax) measures the strength of this period. Microbial ASVs are considered rhythmic ASVs only of PNmax > 0.1 and *p* < 0.05 (detailed in Supplementary Text S4). In order to better observe the dynamics of ASV abundance in response to tidal fluctuations, cluster heatmap was constructed with Centered Logarithm Ratio values (CLR, adding a pseudocount of 1) of rhythmic ASVs. Random forest analyses were performed using the ‘randomForest’ package with default parameters to predict physicochemical variables using rhythmic ASVs. Stratified random sampling was used to select 70% samples to be used as the training dataset and the model were then validated on the remaining 30% samples (detailed in supplementary Text S5). Furthermore, the rhythmic patterns of physicochemical factors were also characterized by LSP analysis with same parameters. The distance-decay relationships (DDRs) [43] of archaeal and bacterial communities in groundwater groups were measured based on taxonomic diversity by linear regression between log-transformed community similarity (Bray-Curtis distance) and geographic distances.

### 2.3. Community assembly processes of tidal creek water microbiota

A recently proposed framework ‘iCAMP’ [22] was used to quantify the contribution of different ecological processes to community assembly based on phylogenetic bin-based null model analysis. The observed taxa were first divided into different ‘bins’ according to a phylogenetic signal threshold (*ds* = 0.2) within the phylogenetic tree and the relative importance of heterogeneous selection, homogeneous selection, homogenizing dispersal, dispersal limitation, and drift in individual ‘bins’ was calculated to evaluate the dominated ‘bins’ and dominated taxa contributing to the ecological processes. In brief, 37294 observed ASVs were divided into 687 phylogenetic bins (renamed to Bin1, Bin2, *etc.*). Phylogenetic tree of 300 most abundant Bins (89.6-97.9% of total contribution to community assembly) and the relative importance of ecological processes in each Bin were displayed by an online tool ‘iTOL’ [44]. Furthermore, all 687 Bins were grouped based on their top member lineage at class level, named ‘Bin-class’. Contributions of the Top 15 and other Bin-class to dominant ecological processes among groups were visualized by R package ‘pheatmap’. In addition, Mantel tests with constrained permutations were performed to reveal the correlations between individual processes and various environmental factors and the scripts are publicly available from GitHub of ‘iCAMP’ (https://github.com/DaliangNing/iCAMP1).

### 2.4. Checkerboard score and co-occurrence network analysis

Checkerboard score (C-score) model was used as the metric of species segregation and its variance (C_var_-score) was used as the metric of both species segregation and aggregation in the intertidal water microbiota [45]. The higher the observed standardized effect (SES) values of C-score, the greater degree of species segregation relative to a random distribution. The C-score tests were performed using the scripts of package MbioAssy1.0 (https://github.com/emblab-westlake/MbioAssy1.0) [31] and more information can be found in Text S6. Co-occurrence network and global network properties were generated using the Molecular Ecological Network Analysis Pipeline [46, 47] and visualized with Gephi v0.9.2. Six networks were constructed based on abundance profiles of ASVs that present in at least half of the samples of each sample group. The resulting correlation matrix (Pearson correlation coefficient) was analyzed with the random matrix theory- (RMT-) based approach to determine the correlation threshold. Moreover, 1000 corresponding Erdös-Réyni random networks, the small-world coefficient (σ) and the keystone nodes were generated for each network and more details are described in Text S7.

## 3. Results

### 3.1. Microbiota diversity and structural variations in intertidal water continuum

We first investigated the diversity and structural variations of prokaryotic (i.e., archaeal and bacterial) communities in intertidal groundwater and surface water. The total concentration of water prokaryotic biomass, as measured by the copy number of 16S rRNA gene, varied from 3.96 × 10^4^ to 6.36 × 10^6^ copies/mL (Table S3), with significantly higher median concentration (Wilcoxon rank-sum test, *p* < 0.05, Fig. S2) in the groundwater or porewater (1.94 × 10^6^ copies/mL) than the surface water (3.61 × 10^5^ copies/mL), as well as in the surface water during the neap tide (5.68 × 10^5^ copies/mL) than the spring tide (3.07 × 10^5^ copies/mL). Tide-driven prokaryotic diversity and structural variations in intertidal groundwater and surface water microbiota were also assessed comparatively. Based on ACE and Shannon indices (Wilcoxon rank-sum test, *p* <0.05, Fig. 1B), archaeal alpha-diversity in the natural tidal creek groundwater (NTG) was significantly higher than (NTS-Flood and NTS-Spring) or equal to (NTS-Ebb and NTS-Neap) that in the surface water, whereas bacterial alpha-diversity showed a reversed trend, i.e., lower in the NTG than the NTS-Ebb, NTS-Neap and NTS-Spring, and equal to the NTS-Flood. Comparatively, human-influenced creek groundwater (HTG) harbored significantly higher prokaryotic alpha-diversity than not only the groundwater (NTG) but also the surface water (NTS) of the natural creek.

When weighing relative abundance, archaeal communities were significantly more abundant in the human-influenced creek groundwater (HTG: 1.76%-49.66%) than the natural one (NTG: 0.56%-14.25%) (Wilcoxon rank-sum test, *p* = 0.02, Table S4), while they were more enriched in the downstream than upstream groundwater of both creeks (Fig. S3). Overall, archaeal communities in both groundwater and surface water were mainly comprised of phylum *Nanoarchaeota* (12.0-76.4%), *Crenarchaeota* (6.2-67.0%), *Thermoplasmatota* (0.3-9.3%), and *Halobacterota* (0.3-53.0%), which showed remarkably dynamics of relative abundance in broad ranges and across sampling groups (Fig. 1C). For example, family *Woesearchaeales* and SCGC AAA011-D5 were the most abundant Archaea in NTG and NTS-Ebb, whereas family *Nitrosopumilaceae* and Marine Group II (typical seawater archaea) were most abundant in NTS-Flood (Fig. S4). Notably, up to 9.8% of unclassified archaea were detected in groundwater, suggesting the prevalence of numerous yet-to-be-characterized ‘archaeal dark matter’ in the coastal zone. Moreover, we observed weak but statistically significant structural differences (PERMANOVA, R^2^ = 0.14, *p* = 0.02, Table S5) in archaeal communities between HTG and NTG (Fig. 2A), as well as higher community dissimilarities within HTG (Wilcoxon rank-sum test, *p* < 0.05). Dynamics of archaeal community structure in the surface water were strongly related to tidal cycles, with similar archaeal structure observed among NTG and NTS-Ebb (R^2^ = 0.06, *p* = 0.202) and significant variations between NTS-Ebb and NTS-Flood (R^2^ = 0.60, *p* < 0.001), which was further confirmed by NMDS ordinations in Fig. 2A. This result well suggested the microbial transport through SGD: i) the transport of groundwater microbes to surface water via groundwater discharge at ebb tide and ii) the transport of coastal microbes to intertidal surface water via seawater intrusion at flood tide.

**Figure 2.**
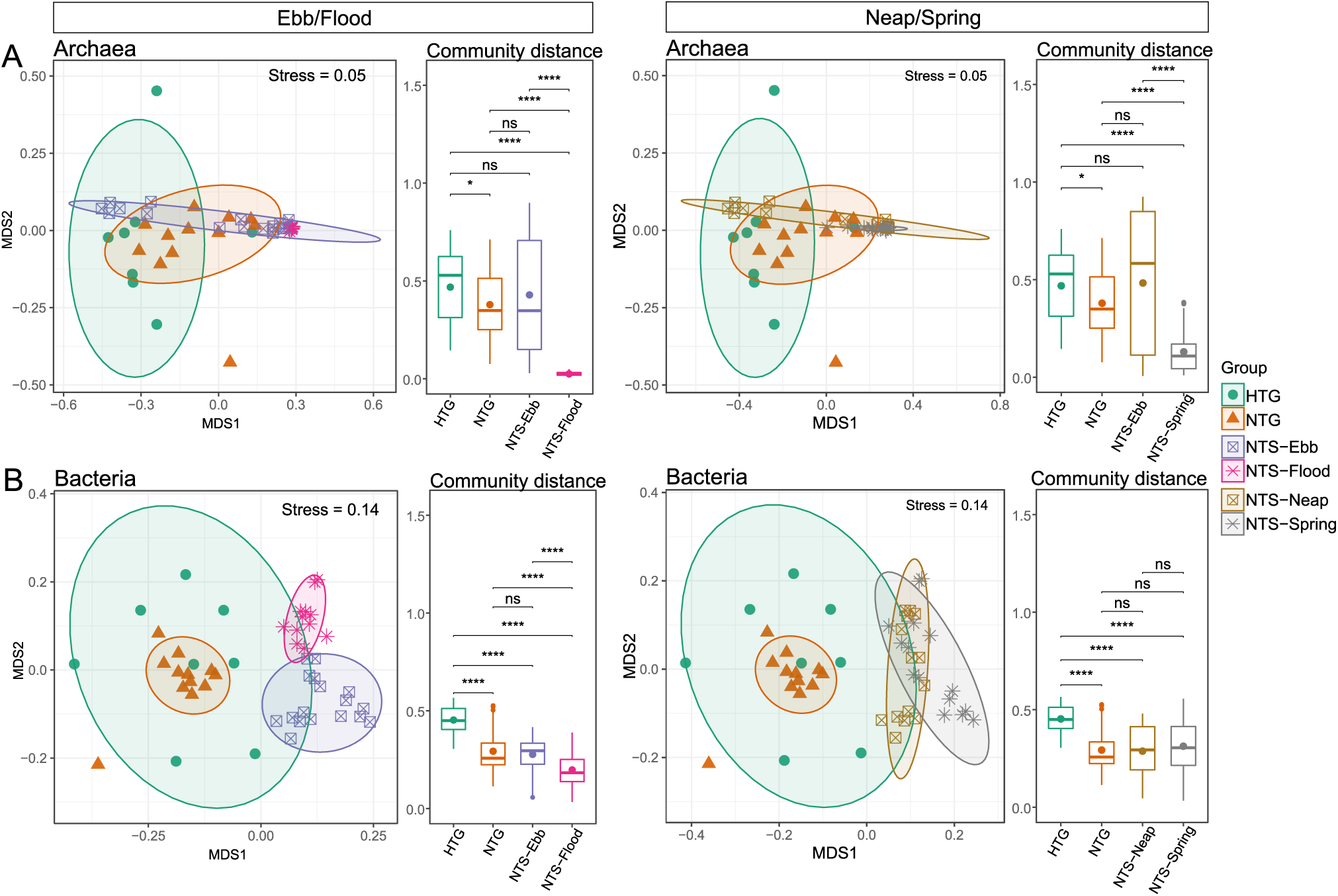
The structural variations of microbial communities within and between tidal creek groundwater and surface water. The non-multidimensional scaling (NMDS) ordination based on weighted UniFrac distance is employed to characterize the compositional variations of archaeal (A) and bacterial (B) communities in HTG, NTG, NTS-Ebb/Flood (left graph) or NTS-Neap/Spring (right graph). The archaeal and bacterial community distance across groups based on weighted UniFrac distance is displayed (right panel) and statistically significant differences across groups are calculated based on wilcoxon rank-sum test (\*\*\*\**p* < 0.0001, \*\*\**p* < 0.001, \*\**p* < 0.01, \**p* < 0.05, ns-non significant).

Echoing archaeal communities, bacterial communities showed significant structural differences between HTG and NTG (R^2^ = 0.16, *p* = 0.002), as well as lower community dissimilarities within the NTG (Fig. 2B). The overall groundwater bacterial communities were dominated by *Proteobacteria* (19.0-82.7%), *Bacteroidota* (1.8-40.7%) and *Campilobacterota* (1.5-39.7%), while the surface water communities were dominated by *Proteobacteria* (28.8-71.5%), *Bacteroidota* (5.5-21.4%), *Cyanobacteria* (1.2-39.7%), *Actinobacteriota* (6.8-22.5%) and *Planctomycetota* (0.8-22.5%) (Fig. 1D). Unlike archaea, bacterial communities were structurally different among NTG, NTS-Ebb, and NTS-Flood (R^2^ = 0.46-0.58, *p* < 0.01, Table S5), as illustrated by their clearly separate clusters in the NMDS diagram (Fig. 2B). Besides, PERMANOVA analysis also showed significant community structural differences between NTS-Neap and NTS-Spring (R^2^ = 0.15, *p* < 0.01).

### 3.2. Microbiota rhythm and spatial dynamic patterns in intertidal surface water

To further resolve the high-resolution temporal dynamics of surface water microbiota, we performed NMDS analysis within the surface water samples (n = 29) to investigate the time-series community structural dynamics at hourly intervals during two whole tidal cycles including neap tide (August 12^th^) and spring tide (August 6^th^-7^th^). Complying with our hypothesis that periodic hydrodynamic disturbance should generate microbiota rhythm patterns, we observed four clusters of surface water samples separated by ebb-flood tide and neap-spring tide with a circular trajectory (Fig. 3A), indicating rhythm patterns of microbiota structure remarkably in the surface water following daily (ebb-flood) and monthly (neap-spring) tidal cycles. The intriguing circadian rhythm patterns of microbiota dynamics were superimposed by highly synchronized and complex bacterial community dynamics and contrastingly weak (but abrupt) and simpler archaeal community dynamics (Fig. S5).

**Figure 3.**
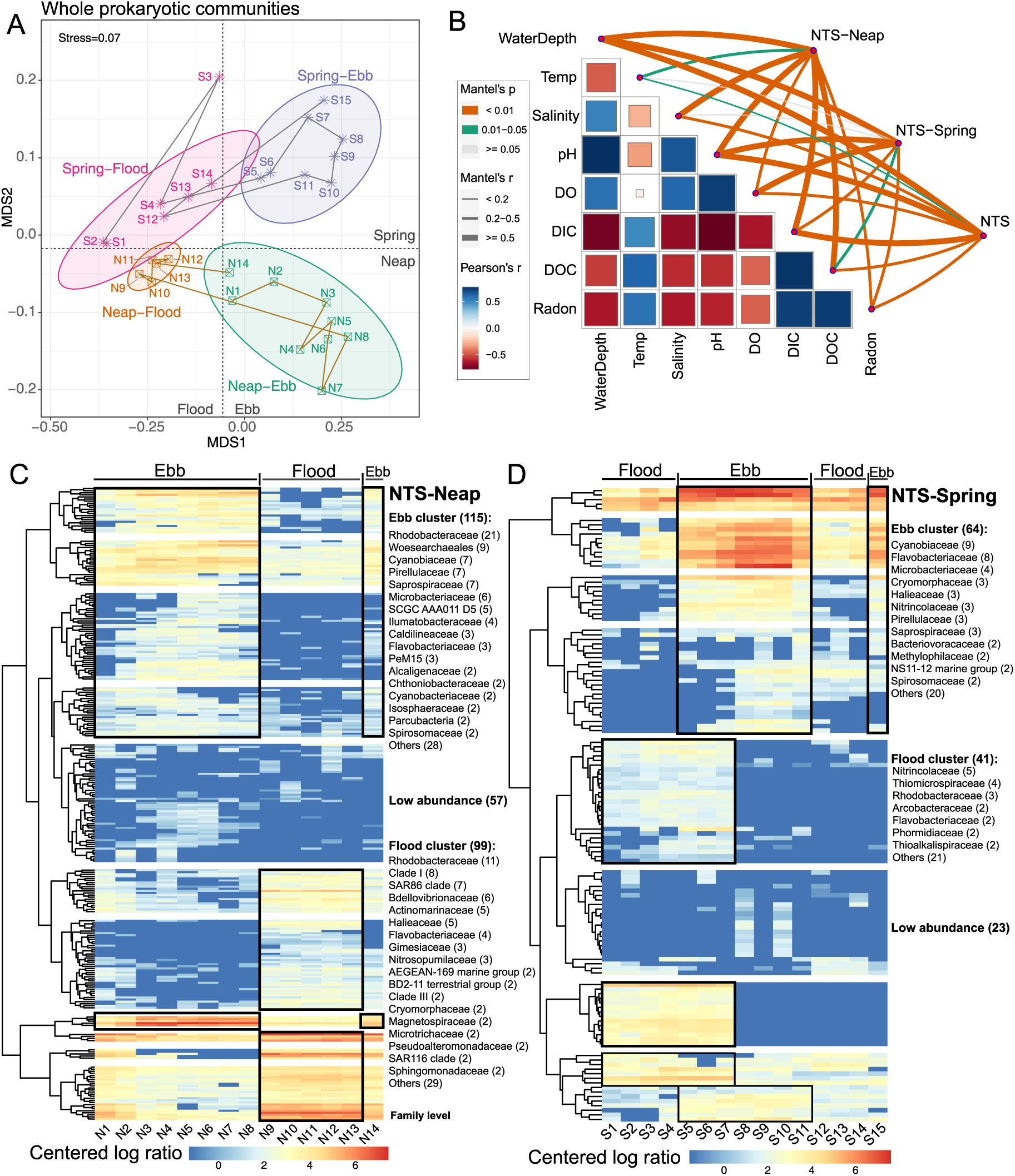
Rhythmicity of microbiota in tidal creek surface water. (A) The NMDS ordination of prokaryotic (archaeal and bacterial) communities in tidal creek surface water is displayed and trajectories show sampling order of surface water during neap tide (N1-N14 with brown lines) and spring tide (S1-S15 with gray lines). The four ellipses mark surface water samples collected during the neap-ebb, neap-flood, spring-ebb and spring-flood tide, respectively. (B) Pairwise comparisons of environmental factors are shown with a color gradient denoting Pearson’s correlation. Microbiota composition in NTS-Neap, NTS-Spring and NTS groups was related to each factor by partial mantel tests. Edge width corresponds to mantel’s r statistic for the corresponding distance correlations and edge color denotes the statistical significance based on 9,999 permutations. (C) Cluster heatmap plot of centered logarithm ratio abundances of 271 rhythmic ASVs identified in the NTS-Neap group. (D) Cluster heatmap plot of 128 rhythmic ASVs identified in the NTS-Spring group. The black rectangles (cluster) mark the ASVs showing higher abundance during ebb tide or flood tide and the number of ASVs classified to each Family for ebb cluster and flood cluster was annotated in the graph.

Further, we correlated surface water microbiota (NTS-Neap, NTS-Spring, and NTS) to eight environmental factors, i.e., water depth, temperature, salinity, pH, DO, DIC, DOC, and ^222^Rn, to verify the aforementioned microbiota rhythm patterns (Fig. 3B). These factors are generally regarded as indicators of tidal cycles [2], and seven of them (except for DOC) were identified to display significant rhythm patterns with tidal cycles (PNmax = 0.34-0.77 and P-value < 0.01, Table S6). The results showed that seven factors, except for temperature, exhibited significant correlations with surface water microbiota during neap tide, spring tide or both, supporting the rhythm patterns of microbiota structure in tidal creek surface water.

In order to test if each microbial ASV exhibited significantly rhythmic pattern (identified as rhythmic ASVs) during tidal time series, that is, their abundance exhibited periodic fluctuations, we performed Lomb Scargle periodogram (LSP) test in NTS-Neap and NTS-Spring, respectively (see Methods and Supplementary Text S4). We identified 271 rhythmic ASVs in NTS-Neap that spanned rhythmic period of 1.9 to 13.0 hours (Fig. S6 and Table S7), accounting for 40.5% to 72.3% of total relative abundance. And 128 rhythmic ASVs (2.0 to 14.0 hours of periods and 6.8% to 47.5% of abundance) were identified from 1686 ASVs in NTS-Spring. Most of the rhythmic period of rhythmic ASVs (13 and 14 h, 335 out of 399) is aligned with the tidal cycles (∼12 h) of the sampling site. Rhythmic prokaryotic subcommunities (i.e., rhythmic ASVs in NTS-Neap and NTS-Spring) exhibited a highly consistent rhythm pattern with the whole communities, whereas weaker rhythmic pattern was observed in the non-rhythmic subcommunities (Fig. S7A), as suggested by the lower variance explained by the first and second axes based on PCoA analysis (weighted-UniFrac distance, 66.2% and 17.1% in rhythmic subcommunities, 36.7% and 23.5% in non-rhythmic subcommunities) (Fig. S7B). This suggested that the rhythmic patterns observed in whole prokaryotic communities were mainly shaped by the numerous and abundant rhythmic subcommunities.

Based on clustering heatmap plot for centered logarithm ratio abundances of 271 rhythmic ASVs from LSP test in NTS-Neap (Fig. 3C), 115 ASVs showed higher abundance at ebb tide (ebb cluster) and 99 ASVs exhibited high abundance at flood tide (flood cluster), whereas the remaining 57 ASVs with low abundance were detected in only a few time points and their rhythmic characteristics were not clear (Table S7). For example, family *Rhodobacteraceae* (21), *Woesearchaeales* (9), and *Cyanobiaceae* (7) dominated in ebb cluster and Clade I (8) and SAR86 clade (7) dominated in flood cluster. For clustering analysis of 128 rhythmic ASVs in NTS-Spring (Fig. 3D), 64 ASVs showed higher abundance at ebb tide. Among them, family *Cyanobiaceae* (9), *Flavobacteriaceae* (8) and *Microbacteriaceae* (4) showed high abundance at ebb tide in both NTS-Neap and NTS-Spring. Forty-one ASVs displayed higher abundance at flood tide in NTS-Spring, for example, *Nitrincolaceae* (5) and *Thiomicrospiraceae* (4), all of which exhibited higher abundance in the early ebb tide after flood tide (Fig. 3D). Remaining 23 ASVs showed low abundance across samples. Overall, the union set of 157 rhythmic ASVs, which obtained by combining rhythmic ASVs of ebb clusters in NTS-Neap (115) and NTS-Spring (64), represented unique and abundant microbes in intertidal groundwater (Table S7). In turn, the other union set of 141 rhythmic ASVs showing higher signals at flood tide (flood cluster) represented unique microbes in coastal seawater. For the rhythmic ASVs at genus level with mean relative abundance greater than 0.1% in ebb-tide and flood-tide groups, we selected 23 rhythmic genera with more than five-fold abundance difference as microbial indicators for intertidal groundwater discharge (e.g., SCGC AAA011-D5) and seawater intrusion (e.g., *Candidatus* Nitrosopumilus), respectively (Table S8). To validate that these rhythmic ASVs were useful indicators of water physiochemistry, eight random forest regression models were constructed to predict each physicochemical variable. Linear regression models comparing the predicted to actual variables showed stronger correlations (adjusted R^2^ = 0.80-0.99, Fig. S8) and regression slopes closer to 1 (0.56-1.15, which represents a perfect prediction), except for temperature and DO, indicating high predictive accuracy and great differentiation between ebb-tide and flood-tide surface water samples.

Besides temporal dynamics of tidal creek water microbiota, we also measured distance-decay relationships (DDRs) based on Bray-Curtis dissimilarity of community structure to understand the spatial microbiota turnovers in intertidal groundwater. Both archaeal and bacterial communities showed significant DDR patterns (P < 0.05), i.e., decreased community similarity with increased geographical distance, whether for the HTG or the NTG (Fig. S9). Moreover, bacterial communities exhibited stronger DDR patterns in the NTG (slope = −0.3443) than HTG (slope = −0.1926), while archaeal communities showed limited differences in their decay patterns between the creeks, revealing that human influence in the coastal zone may reduce the spatial heterogenicity of Bacteria which are overall more sensitive to human influence than Archaea. In particular, archaeal communities showed a far steeper slope (−0.99/−1.09 vs. −0.19/−0.34) and higher fitness value (0.43/0.29 vs. 0.23/0.18) of DDRs than bacterial communities, indicating an overall stronger turnover rate and higher spatial heterogenicity of Archaea than Bacteria in the intertidal groundwater.

### 3.3. Microbiota assembly processes and mechanisms in intertidal groundwater and surface water

To explore mechanisms underpinning spatiotemporal dynamics and patterns in the intertidal GW-SW continuum, the relative contribution of five ecological processes (see methods) to microbiota assembly was quantitatively evaluated. Overall, homogeneous selection (38.5%-69.3%), dispersal limitation (5.2%-49.0%) and drift (8.8%-22.5%) were the dominant processes that together explained 96.3% to 98.5% of community assemblage (Fig. 4A and Table S9), and therefore the subsequent analysis was focused on them. The results showed that homogeneous selection (58.6%-69.3%) predominated in the microbiota assembly of natural GW-SW continuum, including NTG and NTS. And the contribution of dispersal limitation markedly decreased in NTS-Flood (5.2%) and NTS-Spring (7.1%). Compared to the NTG, the importance of dispersal limitation dramatically increased and predominated in the human-influenced groundwater (49.0% in HTG), accompanying decreased homogeneous selection (38.5%) and drift (8.8%).

**Figure 4.**
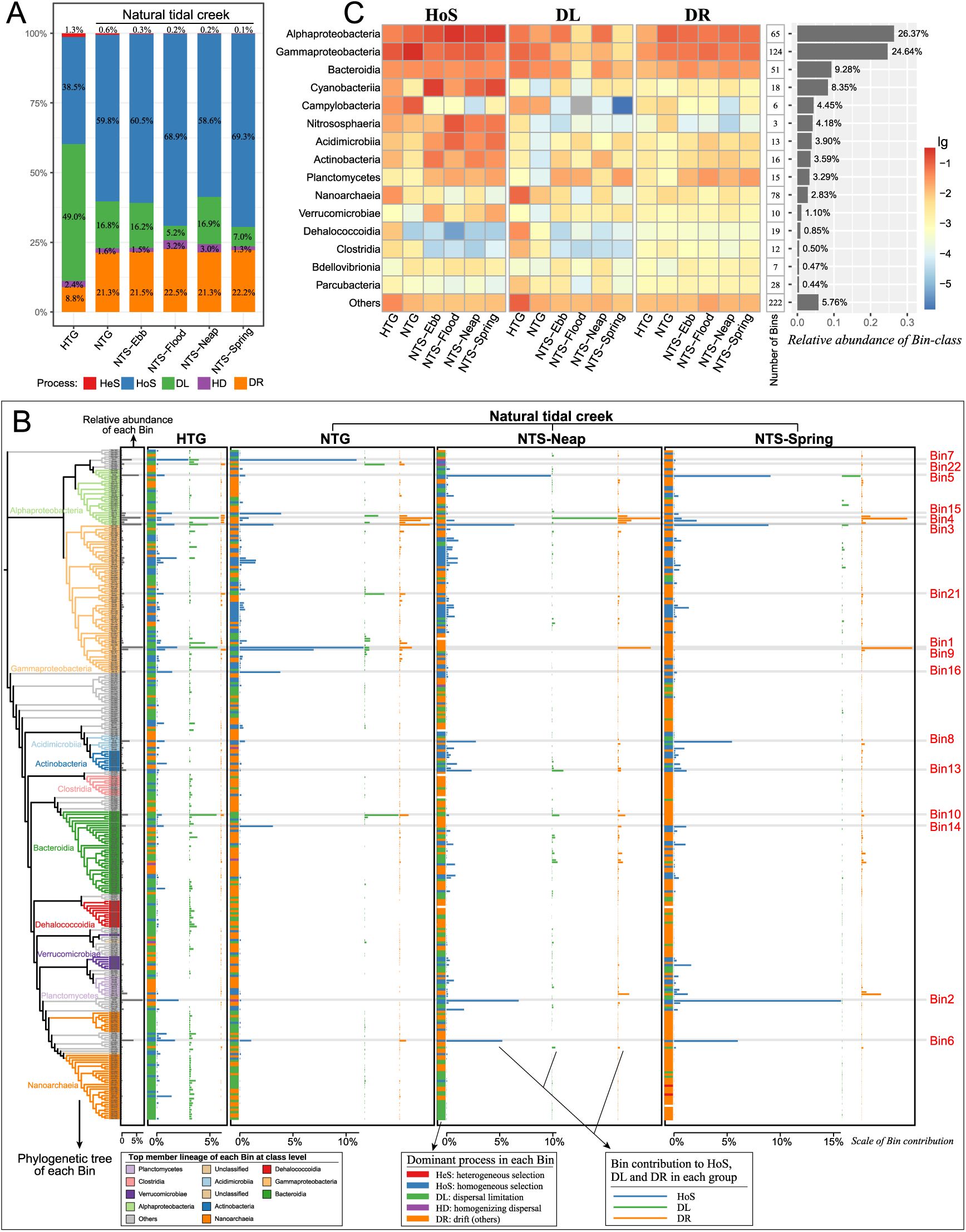
Community assembly processes across different phylogenetic groups. (A) Relative importance of five ecological processes across groups, including heterogeneous selection (HeS), homogeneous selection (HoS), dispersal limitation (DL), homogenizing dispersal (HD), and drift (DR). (B) Phylogenetic tree of top member lineage in each Bin was displayed at class-level. Relative abundance of each Bin was displayed in first rectangle. Dominant processes in each Bin (colored rectangle) and Bin contribution (stacked bars) to major ecological processes (HoS, DL and DR) in HTG, NTG, NTS-Neap and NTS-Spring were displayed in 2-5 annulus. Top Bins with higher contribution to ecological processes were marked by gray strip. Only the top300 Bins were shown in this figure, accounting for a relative abundance of 96.1% of all Bins. (C) Bin-class (top15 and others) contributions to the major ecological processes (HoS, DL, and DR) were displayed in the left heatmap. Sum of the relative abundance of all Bins in each Bin-class was displayed in right bar chart. The vertical numbers at the right of the heatmap showed the number of Bins classified at class-level.

Further, the above-mentioned relative importance of each ecological process was quantified down to the refined level of individual phylogenetic lineages (Bins) defined in iCAMP (Table S11) [22]. Our results revealed that homogeneous selection was dominated in 11.4% to 13.0% of 687 Bins (Table S10), accounting for a summed abundance of 49.3% to 54.1%, while dispersal limitation dominated in 26.1% to 71.0% Bins (abundance: 2.9%-39.2%) and drift dominated in 16.6% to 59.0% Bins (abundance: 10.5%-43.5%), revealing the prevalence and dominance of homogeneous selection in a few species with disproportionally higher abundance in the intertidal GW-SW continuum. By exhibiting the contribution of each Bin, we found that the contribution of a few high-abundance Bins (e.g., the sixteen Bins marked in Fig. 4B) was remarkably higher than that of others and thus cumulatively accounted for the major contribution in the microbiota assembly. For example, the most abundant Bin1 (7.3%) contributed up to 11.6% of homogeneous selection in NTG. The results of grouping Bins at class level, i.e., 104 Bin-class (Table S12), also showed the major contribution of abundant Bin-class in the natural tidal creek (Fig. 4C). For example, the most abundant Bin-class *Alphaproteobacteria*, *Gammaproteobacteria* and *Bacteroidia* contributed most to both homogeneous selection, dispersal limitation and drift in the natural GW-SW continuum. Comparatively, the enhanced dispersal limitation in the HTG was attributed to the jointly increased contribution of high and low abundance Bins, which was confirmed by the higher contribution of Bin-class *Nanoarchaeia* (8.5%), *Dehalococcoidia* (3.6%), and *Clostridia* (1.4%) (Fig. 4C). Overall, these results indicated that high-abundance microbial lineages contributed most to homogeneous selection and drift in the natural intertidal GW-SW continuum, whereas the accumulated contributions of both abundant and rare lineages resulted in the enhanced dispersal limitation in the human-influenced groundwater.

### 3.4. Environmental factors influencing the ecological processes of microbiota assembly

Correlations between changes in environmental factors and dominant ecological processes among paired samples in each group were further determined with Mantel test (*p* < 0.05) to identify the major drivers of ecological processes. Overall, homogeneous selection and dispersal limitation showed statistically similar correlation patterns (Mantel test for p-value correlation matrix: r = 0.70, *p* < 0.05) with various environmental factors, while weaker and less frequent correlations were detected between drift and environmental factors (Fig. 5 and Table S13). Both homogeneous selection and dispersal limitation showed significant correlations with salinity, pH, DO, DIC and DOC in NTS-Neap and NTS-Spring. Considering the fluctuating environmental factors with tidal flows and their significant inter-correlations (Fig. S10), significant correlations between these factors and dispersal limitation may indicate the influence of GW-SW mixing on the temporal microbiota turnover in intertidal surface water. Notably, the environmental factors associated with homogeneous selection were quite different in the HTG (DIC and DOC) and NTG (salinity, pH and DO), and so was dispersal limitation and drift, suggesting a potential effect of carbon nutrients on the microbiota assembly in the human-influenced groundwater.

**Figure 5.**
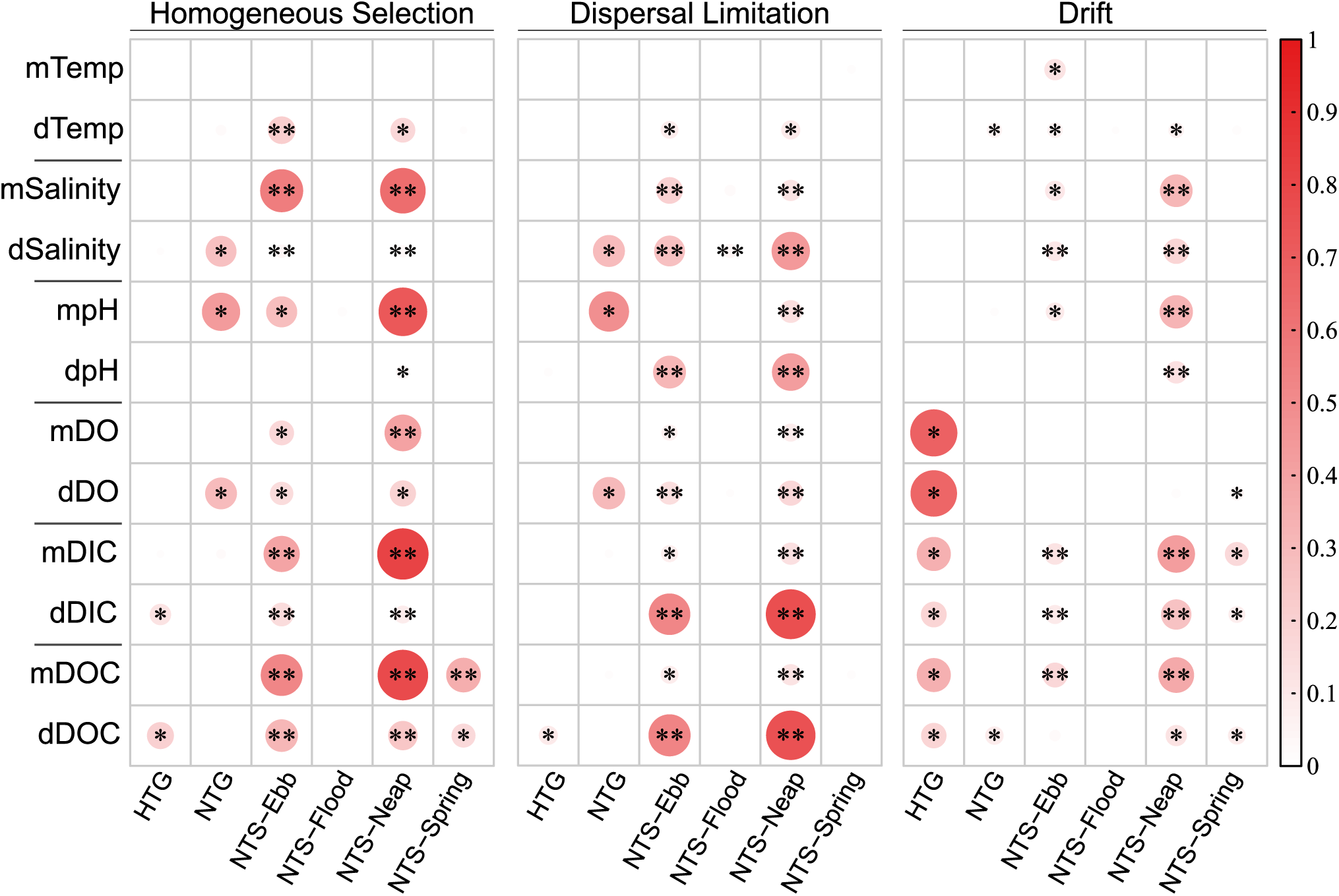
Effects of environmental factors on major ecological processes. Heatmap plot of correlations between environmental factors and dominant ecological processes based on mantel test. The correlation was determined based on the difference (with a ‘d’ before the factors) or the mean (with a ‘m’ before the factors) of a factor between each pair of samples. Significance was expressed as \*\**p* < 0.01, \**p* < 0.05.

### 3.5. Deterministic microbial segregation and co-occurrence patterns of tidal creek water microbiota

Once stochastic and deterministic processes are demonstrated to co-drive the microbiota assembly in the intertidal GW-SW continuum, we applied C-score and co-occurrence network analyses to explore the contribution of microbial interactions (e.g., competition and mutualism) [22] on the observed assembly patterns, which are included in selection process but could not parse out by the null model analysis. Based on positive values of standardized effect size (SES) of C-score and their ranks (Fig. 6A), the intertidal water microbiota exhibited significant and stronger segregated distributions during Neap than Spring tide (51.2 vs 21.8), during Ebb than Flood (21.8 vs 13.2), and in HTG than NTG (13.9 vs 5.3), compared with those expected under ‘random’ null hypothesis (Table S14). Moreover, the SES values of both C-score (5.3) and C_var_-score (3.9) were the lowest in NTG (5.3 and 3.9), while their values were much higher in HTG (13.9 and 11.7), revealing an overall lower degree of both species segregation and aggregation (e.g., resulted from competition and cooperation) in the natural than the human-influenced groundwater.

**Figure 6.**
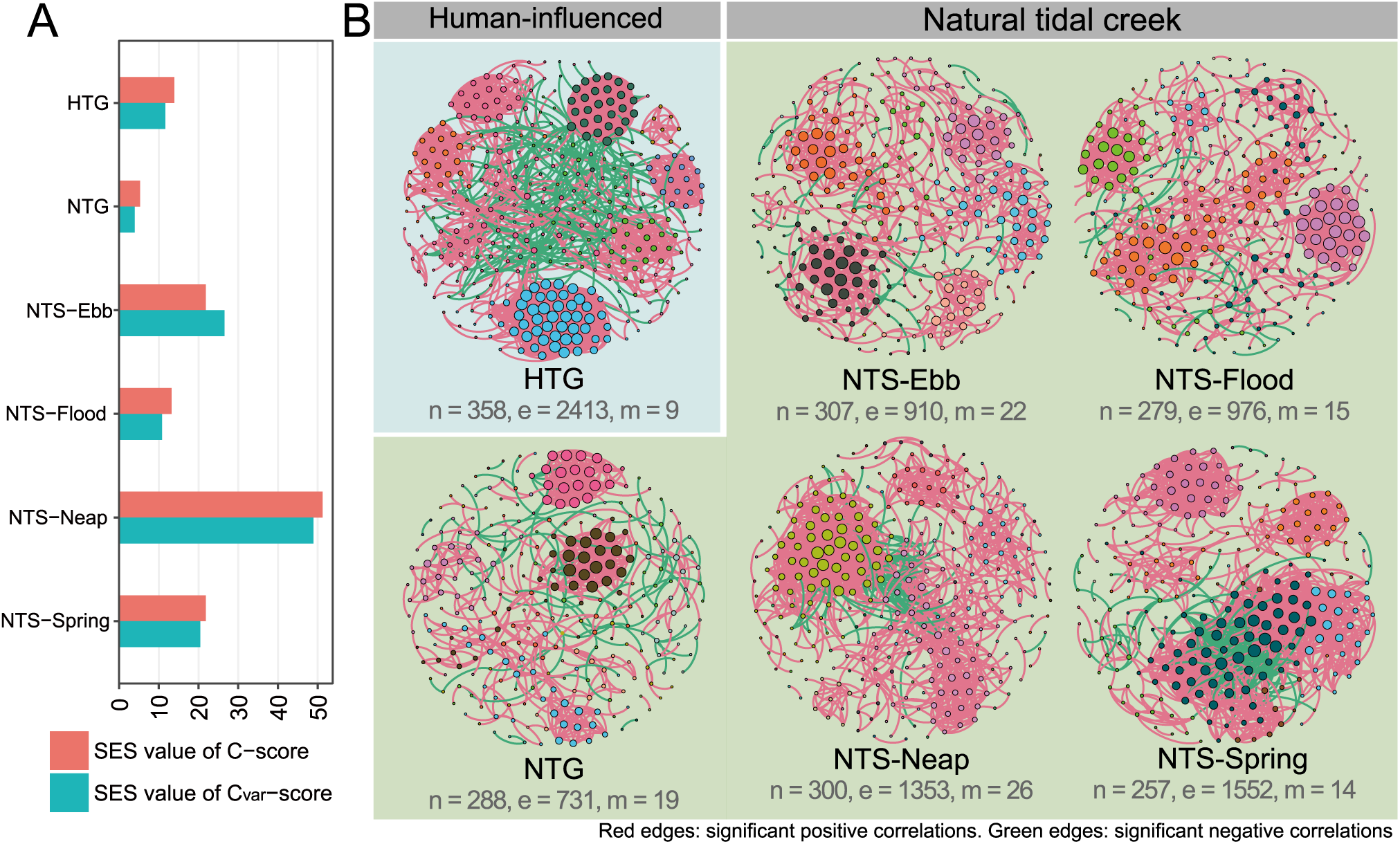
Microbial segregation and co-occurrence patterns of tidal creek water microbiota. (A) Standardized effect sizes (SES) values of C-score and C_var_-score for intertidal groundwater and surface water microbiota over six groups. Higher observed SES values of C-score suggest greater degrees of species segregation than would be expected by chance. Higher SES values of C_var_-score indicate greater degrees of both species segregation and aggregation. All *P*-values are < 0.0001 (detailed in Text S6 and Table S14). (B) Microbial co-occurrence networks in groundwater and surface water groups. n: number of nodes; e: number of edges; m: number of modules. Nodes in each network are colored by modularity. Red edges represent significant positive correlations and green edges represent significant negative correlations. More details are described in Text S7 and Table S15.

Because C-score analysis revealed significant roles of both cooperative (e.g., synergistic) and competitive (e.g., antagonistic) microbial interactions in microbiota assembly [48], we further explored microbial co-occurrence patterns by establishing six networks for intertidal groundwater or surface water microbiota (Fig. 6B). Topological analysis showed that the networks displayed significantly different topological structures from their corresponding random networks of identical size. They all had highly modular structures (modularity > 0.4) and showed ‘small-world’ properties (σ > 1) with coefficients between 1.93 to 6.19 (Table S15), revealing non-random network structures and assembly patterns that were unlikely to appear due to chance. In the natural intertidal GW-SW continuum, intriguingly, a higher percentage of rhythmic ASVs was observed in each surface water network (61.5%-73.7%) than in the groundwater network (44.8%, Table S15). Meanwhile, seven of the eight keystone taxa (5 module hubs and 3 connectors) identified in the surface water network were also rhythmic ASVs (Table S16). The key role of rhythmic ASVs observed in the surface water networks suggested the impact of tidal fluctuations on microbial interactions in intertidal surface water.

The anthropogenic influence on co-occurrence patterns of groundwater microbiota was comparatively explored between HTG and NTG (Fig. 6B). The number of nodes (358), edges (2413), graph density (0.038), and average clustering coefficient (0.491) of HTG network were all remarkably higher than that of NTG network (288, 713, 0.018 and 0.308, respectively), suggesting higher frequencies of species co-occurrence associations in the human-influenced than natural groundwater. Moreover, mantel tests showed that both temperature and DIC/DOC correlated positively (though not always significant) with almost all modules in HTG network (*P* < 0.05, Fig. S11), whereas none of these parameters correlated with modules in NTG network, suggesting they played a potential essential role in shaping the modularized structures in HTG network representing discrete ecological niches in the human-influenced tide creek. Ten keystone ASVs were observed in HTG network and were all identified as ‘connector’, but only two keystone ASVs (one ‘module hub’ and one ‘connector’) were identified in NTG network (Table S16 and Fig. S12).

## 4. Discussion

Tidal hydrodynamics drive the groundwater-seawater exchange and shifts in microbiota structure in the intertidal zone. However, the periodic tidal fluctuations and complex geological environment make it difficult to accurately resolve the temporal dynamics of microbiota structure based on a few discrete time-series samples in this intertidal groundwater-surface water continuum. Through hourly time-series microbiota profiling and modeling, this study provides first insights into the microbiota rhythmic patterns in response to periodic tidal fluctuations in intertidal surface water of Yancheng, Jiangsu Province, China, and identifies homogeneous selection as the key process driving microbiota structuring and assembly in the intertidal GW-SW continuum. Furthermore, by comparing the groundwater microbiota of a tidal creek affected by anthropogenically contaminated river inputs with the natural creek, we found that shifts in prokaryotic biodiversity and community structure were accompanied by enhanced dispersal limitation, drift and microbial interactions but weakened homogeneous selection.

### 4.1. Tide cycles drive periodic transport and rhythm patterns of microbiota along the natural intertidal water continuum

The high-resolution temporal dynamics of microbiota structure in tidal creek surface water were followed during whole tidal cycles. Our results revealed that tidal fluctuations shaped the daily (i.e., ebb-flood) and monthly (i.e., neap-spring) dynamics of microbiota structure. Unlike archaea transport mainly via groundwater discharge during ebb tide, potential multi-source transport mechanisms other than groundwater discharge may account for the unique bacterial community structure of ebb-tide surface water that differs from the intertidal groundwater. For example, bacterial biota might also originate from the release of particle-attached bacteria via particle resuspension from surface sediments and flushing of the surficial sediment bacteria [49]. The likelihood of such events in our study is confirmed by the unique higher abundance of family *Cyanobiaceae* (Fig. S4) and genera *Candidatus* Aquiluna, *Roseibacillus*, *NS11-12 marine group* (Fig. S13) in the ebb-tide surface water compared to the intertidal groundwater and flood-tide surface water. The family *Cyanobiaceae* have been reported to prefer to develop on the surface of saltmarsh tidal mudflats [7, 50]. Taken together, tidal fluctuations pose a domain-level community-specific influence on the transport mechanisms of Archaea and Bacteria. This discrepancy may be related to the fact that archaeal prefer to live in the subsurface sediment, while bacteria have a wider niche breadth in the intertidal zone [51].

Especially, based on the hourly time-series sampling of intertidal surface water, we have further found that periodic tidal fluctuations on a daily and monthly basis lead to the rhythm patterns of microbiota structure. The rhythmicity of environmental water microbiota has only been implicated in several long-term studies on the seasonal or circadian reoccurring patterns of marine water community structure [42, 52, 53]. Although the periodic shift of coastal microbiota structure associated with tidal fluctuation has been reported before through observation of a few discrete-time samples [49], our results based on hourly time-series sampling demonstrated that tidal cycles dominated the rhythmic recurrent of not only groundwater-related microbiota during ebb tide but also seawater-related microbiota during flood tide. The rhythmic ASVs considered as microbial indicator of intertidal groundwater discharge (ebb-tide) and coastal seawater (flood-tide) shared very few at the family level (6 of 214 shared in NTS-Neap and 13 of 105 shared in NTS-Spring, Table S7). Even for these shared family, they are classified as different genera. The rhythmic subcommunities suggested that different ecological niches among intertidal groundwater and coastal seawater selected for different microbial populations, which was responsible for the unique but abundant biota observed in ebb-tide and flood-tide, respectively. In construct, the non-rhythmic subcommunities with similar abundance in ebb and flood tide surface water should be adapted to both groundwater and seawater environment, and dispersal with the bidirectional water exchange. This was confirmed by the higher contribution of rhythmic genera to homogeneous selection relative to dispersal limitation in intertidal groundwater (e.g., genera *Cyanobium* PCC-6307 and *Rhodobacteraceae*) (Table S10). Intriguingly, the rhythm pattern during spring tide was weaker (fewer rhythmic ASVs) than during neap tide, probably because the high tide level inundates the saltmarsh and overlying water is discharged together with the groundwater at ebb tide, interfering with the characterization of rhythm pattern. Besides, change of tidal periods in the investigated tidal creek resulting from tidal asymmetry (e.g., decrease in the duration of ebb and increase in the duration of flood) [54], as well as the hysteresis patterns on different time scales (e.g., diel and neap-spring) [35], may explain the occurrence of rhythmic period shorter than tidal cycles.

Furthermore, the random forest models represented high predictive accuracy in physicochemical variables and great differentiation between ebb-tide and flood-tide surface water samples. Given that these variables, e.g., salinity, pH [55] and ^222^Rn [56], were commonly used as environmental tracers to establish intertidal SGD or seawater intrusion in the coastal zone, our results demonstrated the potential of microbiome analysis to enhance monitoring of coastal water quality and pollution sources by detecting rhythmic microbial variances that signify anthropogenic contamination from intertidal groundwater discharge and/or seawater intrusion. For example, the presence of high abundance species such as *Vibrio mimicus* (0.47%) and *Vibrio cholerae* (0.15%) in HTG-1M, which are important pathogens associated with aquaculture [57], suggested potential aquaculture pollution in intertidal groundwater. This finding supports the hypothesis that high abundance of such fecal or aquaculture indicators, detected as ebb-tide rhythmic ASVs in coastal seawater, may indicate coastal anthropogenic pollution resulting from intertidal groundwater discharge. In addition, among the 23 rhythmic genera we selected as microbial indicators, SCGC AAA011-D5 and *Woesearchaeales* belonging to order *Woesearchaeales* were also reported to be predominant (45%-76% of archaea) in intertidal unconfined aquifer but accounted for less than 1% in seawater, whereas genera *Candidatus* Nitrosopumilus and Marine Group II exhibited the opposite trend [58]. This suggested that these rhythmic ASVs and filtered rhythmic genera have a broad applicability in indicating coastal groundwater discharge and seawater intrusion. Further validation can be achieved through larger-scale multi-habitat studies in the future. We propose to leverage the informatics of microbiota to add biologically relevant information to the monitoring and assessment of coastal ecosystem.

### 4.2. Homogeneous selection and microbial interactions co-contribute to the microbiota assembly in the natural intertidal water continuum

Given the tide-driven hydrodynamic disturbance and the accompanying environmental gradients across the temporal and spatial scales in the coastal zone [59], the corresponding ecological processes, i.e., selection and dispersal, are expected to play a key role in the microbiota assembly in the natural tidal creek. Despite microbial dispersal is expected to be enhanced under periodic tidal fluctuations, high levels of dispersal can result in a homogenization of the local and regional species pool, which can lead to an overall short-time increase in stochasticity (during initial formation phases of tidal creeks) followed by an increase in deterministic selection at a regional scale that decreases species diversity [60] (Fig. 1B). Meanwhile, the enhanced homogeneous selection can lead to increased species competition, suggested by the species segregated distributions based on C-score analyses. Compared with the surface water, weakening of water mixing disturbance in intertidal groundwater resulted from the barrier effect of cohesive sediment can account for the relatively lower dispersal rates. We conclude that homogeneous selection and microbial interactions co-contributed to the microbiota assembly patterns in the natural intertidal water continuum.

### 4.3. Anthropogenic influences alter microbiota structure and community assembly patterns in intertidal groundwater

Intertidal zone harbors abundant and diverse microbial environments owing to the land-sea interaction and are extremely sensitive to natural changes and nearshore human activities [50, 61]. Nutrient loading derived from human activities and tidal dynamics is among the critical pressures affecting the intertidal environments [62]. Considering the high spatial heterogeneity of intertidal environment, we selected two geographically adjacent (within 6 km) tidal creeks for comparative analysis of intertidal groundwater microbiota, so as to maintain a basically consistent environmental conditions except for the single variable of anthropogenic disturbance in NT. Our study found that the human-influenced tidal creek exhibited both higher species diversity and community structural dissimilarities of groundwater microbiota than the natural one, whether for archaeal or bacterial communities. This contrasting profile is expected to result from the shift in microbiota structure caused by polluted creek surface water intrusion due to Terrestrial River water input upstream (during ebb) and estuary-water input downstream (during flood) of the creek.

Compared with the natural groundwater, the human-influenced groundwater showed remarkably increased (and dominant) dispersal limitation, decreased homogeneous selection and drift, and stronger microbial co-occurrence patterns. First, the river water input carries abundant nutrients from aquaculture and human activities, which can create additional niches to stimulate the growth of various microorganisms, especially less abundant ones. Such growth could increase species diversity (Fig. 1B) and enhance stochastic processes like birth and death, the latter of which is consistent with the enhanced dispersal limitation (the influence of dispersal limitation acting in concert with drift [23]) and drift (the influence of drift acting alone) in the human-influenced groundwater, as noticed in prior studies on terrestrial freshwater [63] and groundwater [13]. Second, nutrient inputs could weaken environmental selection by providing more resources and result in less-stressed groundwater conditions, partly resulting in decreased homogeneous selection. Nutrient-associated factors dominated selection process in the human-influenced groundwater, suggested by the relationships between homogeneous selection and DIC or DOC (Fig. 5). Higher abundance of nitrogen-metabolizing genera, e.g., nitrifying *Nitrosopumilus*. [64] and denitrifying *Hydrogenophag*a [65, 66], and carbon-metabolizing genera, e.g., *Bathyarchaeia* [67], and *Sphaerochaeta* [68] in the human-influenced than the natural groundwater (Fig. S14) further supported the important effect of nutrient input. Third, exogenous water inputs at ebb tide provided supplement water and therefore weakened tidal fluctuations (i.e., weakened groundwater discharge during ebb tide), leading to enhanced dispersal limitation.

Further, our finding on the important role of stochastic processes (dispersal limitation in concert with drift) in controlling the human-influenced groundwater microbiota is consistent with the prior studies indicating that stochasticity decreased, whereas environmental selection could be stronger in a fluidic ecosystem with a disturbance, in turn, stochasticity increases after adding exogenous nutrients [13]. Moreover, the combined contribution of dispersal and homogeneous selection have also shaped DDR patterns of intertidal groundwater microbiota. However, it seems that nutrient-driven selection processes dampened the effect of dispersal limitation, resulting in decreased bacterial community dissimilarities across geographic distance in the human-influenced groundwater compared with the natural groundwater. Interestingly, unlike bacterial communities, archaeal communities showed little change in DDR patterns in the human-influenced groundwater, potentially due to a narrow niche breadth of subsurface archaea and thus less influenced by anthropogenic activities [51].

Despite its lower importance of homogeneous selection (environmental filtering), the human-influenced groundwater microbiota showed both a higher degree of species segregation and higher frequencies of species co-occurrence associations, indicating stronger microbial interactions. On the one hand, enhanced dispersal limitation and potentially stronger competition (increased species diversity with similar ecological niches) enhanced the species segregation distribution. On the other hand, complex physicochemical gradients (e.g., abundant nutrients, salinity and DOC) [69] created multiple niches that harbor various microbial groups with different metabolic capacities (Fig. S11) and in turn created opportunities for synergistic or other cooperative interactions (mutualistic and commensal metabolism cooperation), which was supported by enrichment of copiotrophic genera *Woesearchaeales* [70] and *Flavobacterium* [71] and more in the human-influenced groundwater (Fig. S14). Meanwhile, more keystone ‘connectors’ enhanced the stability of the community under higher level of microbial cooperation (reduce stability) [72] in the human-influenced groundwater. Overall, the enhanced microbial interaction is one of the essential ecological processes that cannot be ignored when exploring the microbiota assembly in the human-influenced intertidal groundwater.

## 5. Conclusions

By overcoming obstacles of time-series sampling in the salt marsh, this study first unravels microbiota rhythm patterns in response to periodic tidal fluctuations in the intertidal groundwater-surface water continuum where microbiota assembly was found to be dominantly driven by homogeneous selection. Discharge of anthropogenically polluted river water into the tidal creek is demonstrated to remarkably alter the microbiota biodiversity, structure and assembly patterns in the intertidal groundwater via bidirectional water exchange. Our first combined use of rhythm analysis and random forest models suggested the potential to enhance monitoring of coastal water quality and pollution sources by detecting rhythmic microbial variances that signify anthropogenic contamination from intertidal groundwater discharge and/or seawater intrusion. This study also reveals how the anthropogenically polluted river input shift intertidal groundwater microbiota via water exchange, which in turn may affect the microbially-driven biogeochemical cycling in coastal aquifers. Altogether, this study can provide valuable insights into the bio-monitoring and conservation of coastal zone ecosystems.

## Data Availability

The raw sequencing data generated in this study have been deposited in the China National GeneBank DataBase (CNGBdb) with accession number CNP0003642.

## Acknowledgment

This work was supported by the Zhejiang Provincial Natural Science Foundation of China under Grant No. LR22D010001 and the HRHI program 202309010 of Westlake Laboratory of Life Sciences and Biomedicine. We acknowledge the Westlake University HPC Center for computational support. Thank Mr. Duofei Hu and Ms. Shengjie Hu for assistance in field sampling, and Ms. Yisong Xu for laboratory management supports.

## Author Contributions

All authors contributed intellectual input and assistance to this study. F.J. obtained funding and supervised the project. The original concepts were conceived by F.J. and Z.Z. with advice from L.L. Field sampling was carried out by Z.Z. and X.C. Data analysis were done by Z.Z. with assistance from G.Z. The manuscript was prepared by Z.Z. and F.J. and revised by L.L., L.Z. and H.G.

