## Supplemental text and figure for "Hydrodynamic and anthropogenic disturbances co-shape microbiota rhythmicity and community assembly within intertidal groundwater-surface water continuum"

##### Supplementary Methods

**Text S1.** Physiochemical monitoring of water samples

**Text S2.** Real-time quantitative PCR for 16S rRNA gene copies

**Text S3.** 16S rRNA gene sequencing and QIIME2 analysis

**Text S4.** Lomb Scargle periodogram analysis of microbial ASVs and environmental factors

**Text S5.** Using rhythmic prokaryotic subcommunities to predict physicochemical variables

**Text S6**. Standardized Checkerboard Index analysis for intertidal water microbiota

**Text S7.** Co-occurrence network analysis

##### Supplementary Figure

**Fig. S1.** Time series sampling of surface water in the times series station of natural tidal creek (NT) and geochemical observations of intertidal surface water measured in time series station at neap tide and spring tide

**Fig. S2.** Prokaryotic biomass concentration of groundwater and surface water in Saltmarsh Tidal Creek of Yancheng National Nature Reserve, China

**Fig. S3.** Kingdom-level microbial composition in tidal creek groundwater and surface water

**Fig. S4.** The relative abundance of family-level archaeal and bacterial communities.

**Fig. S5.** The NMDS ordination of archaeal and bacterial communities in intertidal surface water based on weighted UniFrac distance

**Fig. S6.** Number of rhythmic and non-rhythmic ASVs with different rhythmic periods

**Fig. S7.** The NMDS and PCoA ordination of rhythmic subcommunities and non-rhythmic subcommunities in intertidal surface water

**Fig. S8.** Predicted physicochemical values based on random forest regression model versus actual values

**Fig. S9.** Distance-decay relationships (DDRs) of archaeal and bacterial communities

**Fig. S10.** Correlations between each pair of environmental factors

**Fig. S11.** Heatmap plots of correlations between environmental factors and microbiota in each module of each co-occurrence network

**Fig. S12.** Putative keystone taxa identified based on the node topology in each network.

**Fig. S13.** Heatmap plots of relative abundance of genus-level prokaryotic communities in natural intertidal groundwater-surface water continuum.

**Fig. S14**. Heatmap plots of relative abundance of genus-level prokaryotic communities in intertidal groundwater

### Supplementary methods

#### Text S1: Physiochemical monitoring of water samples

Water temperature, pH, salinity and dissolved oxygen (DO) of groundwater samples were determined in situ by a multi-parameter sonde (Multi 3430 WTW, Germany). For the time-series observations, temperature, pH, salinity, water depth and DO were measured in situ using an automated datalogger (EXO3 Multiparameter Sonde) at 1-min intervals. $^{222}$Rn was measured in situ by a RAD7 detector. Water samples for dissolved inorganic carbon (DIC) and dissolved organic carbon (DOC) analysis were filtered in situ through 0.45-μm nylon filters. These filtrates were treated with saturated $HgCl_2$ and stored at 4 °C until analyzed using a total organic carbon analyzer in the laboratory (TOC-VCPH, Shimadzu Co. Ltd., Japan). Measurement details and physicochemical data were provided in our earlier publication [1].

#### Text S2: Real-time quantitative PCR for 16S rRNA gene copies

Prokaryotic 16S rRNA gene copies was quantified by real-time quantitative PCR of total DNA in each sample using primes 515F (5'-GTGYCAGCMGCCGCGGTAA-3') and 926R (5'-CCGYCAATTYMTTTRAGTTT-3'), which targets both archaea and bacteria. Each 20 μL reaction mixture volume was consisted of 10 μL SYBR Premix Ex TaqTM (TaKaRa, Japan), 0.5 μL each primer, 1.0 μL template DNA and 8 μL nuclease-free PCR-grade water. The thermal cycle was consisted of a 10 min initial enzyme activation at 95 °C, followed by 40 cycles of denaturation at 95 °C for 30 s, annealing at 60 °C for 30 s and extension at 72 °C for 15 s. A plasmid control containing a cloned and sequenced 16S rRNA gene fragment (1.24E+10 copies/L) was used to generate six-point calibration curves from tenfold dilutions for standard calculation. All qPCR reactions were performed in technical triplicates for each DNA extract, standard, and negative controls using LightCycler® 480 Probes Master (Roche, Basel, Switzerland). The reactions were performed in 96-well plates on a Jena qTOWER$^3$ G (Germany).

#### Text S3: 16S rRNA gene sequencing and QIIME2 analysis

The V4-V5 regions of prokaryotic 16S rRNA genes were amplified using primers 515F (5'-GTGYCAGCMGCCGCGGTAA-3') and 926R (5'-CCGYCAATTYMTTTRAGTTT-3') and sequenced on the Illumina NovaSeq 6000 platform with 250-bp paired-end reads at the Magigene Biotechnology Co. Ltd (Guangzhou, China). The raw sequencing data was processed using QIIME2 pipeline [2] and DADA2 algorithm [3]. First, FastQC are employed to get quality score of the raw reads. Then, paired-end sequences were truncated from bases 19 to 240 and 20 to 220 for forward and reverse sequences, respectively, to meet the requirement of quality score above 25. After quality trimming, merging of paired sequences and removal of chimeric sequences, all reads were subjected to de-noising using the DADA2 pipeline based on 100% sequence similarity with default parameters. Then, a 16S rRNA V4V5 specific classifier was trained with Silva small subunit rRNA sequences (release 138.1 [4]) by q2-feature-classifier plugin in QIIME2. Taxonomic assignment of amplicon sequence variant (ASVs) from DADA2 were performed with the classifier. The ASVs annotated as chloroplasts or mitochondria were excluded for further analysis. The phylogenetic tree was generated (FastTree plugin) from representative sequences after aligned by mafft.

**Text S4: Lomb Scargle periodogram analysis of microbial ASVs and environmental factors**

The Lomb Scargle periodogram (LSP) was used to determine if periodic patterns were present in microbial ASVs according to previous studies testing seasonal and yearly rhythm of marine microbial communities [5, 6]. The LSP analysis was accomplished via the lomb package (version 2.1.0) in the R software [7] and the script was modified from a previous study [5] (https://github.com/adriaaulaICM/bbmo_niche_sea). The LSP determines the spectrum of frequencies composing the dataset. For each ASV, we obtain the density distribution for each of the periods and the peak normalized power (PNmax). The distribution shows which are the most recurrent period and the PNmax measures the strength of this period. In this study, we considered the microbial ASVs as rhythmic ASVs only if PNmax was above 0.1 and $p < 0.05$. The LSP analysis was performed by relative abundance table of ASVs in NTS-Neap and NTS-Spring, respectively, and only ASVs in at least two samples were included. We identified 271 rhythmic ASVs from 1705 ASVs in NTS-Neap and 128 rhythmic ASVs from 1686 ASVs in NTS-Spring (Fig. S6 and Table S7). Then, non-metric multidimensional scaling (NMDS) analysis of rhythmic prokaryotic subcommunities (i.e., all rhythmic ASVs in NTS-Neap and NTS-Spring) and non-rhythmic prokaryotic subcommunities (i.e., all non-rhythmic ASVs) were performed based on weighted UniFrac distance, respectively. Cluster heatmap plot was constructed with Centered Logarithm Ratio values (CLR, adding a pseudocount of 1) of rhythmic ASVs in NTS-Neap and NTS-Spring, respectively. Cluster of rhythmic ASVs showing high abundance at ebb tide (ebb-cluster) or flood tide (flood cluster) surface was marked in the heatmap graph (Fig. 3CD). Finally, these rhythmic ASVs were further classified as potential indicators of intertidal groundwater discharge or coastal seawater according to the detection of higher recurrent signal at ebb tide or flood tide, respectively. In addition, the rhythmicity of environmental factors, including temperature, salinity, pH, DO, DIC, DOC and Radon, were also determined via LSP analysis and the results were showed in Table S6.

Here, as an example, we exhibited the temporal abundance trends of top 5 most abundant rhythmic and non-rhythmic ASVs in NTS-Neap in the figure below. As can be seen from the figure, the abundance of rhythmic ASVs displayed rhythmic patterns in response to tidal fluctuations (A), that is, they exhibited difference abundances during ebb tide and flood tide and varied with tidal cycles. Specifically, the abundance of rhythmic ASVs increased or decreased significantly at the early time point of ebb tide (e.g., N1, N2 and N14), and showed the opposite trend at the early stage of flood tide (e.g., N9). In contrast, the abundance of non-rhythmic ASVs did not exhibit such periodic changes, but showed multiple peaks and valleys and or stable abundance (B).

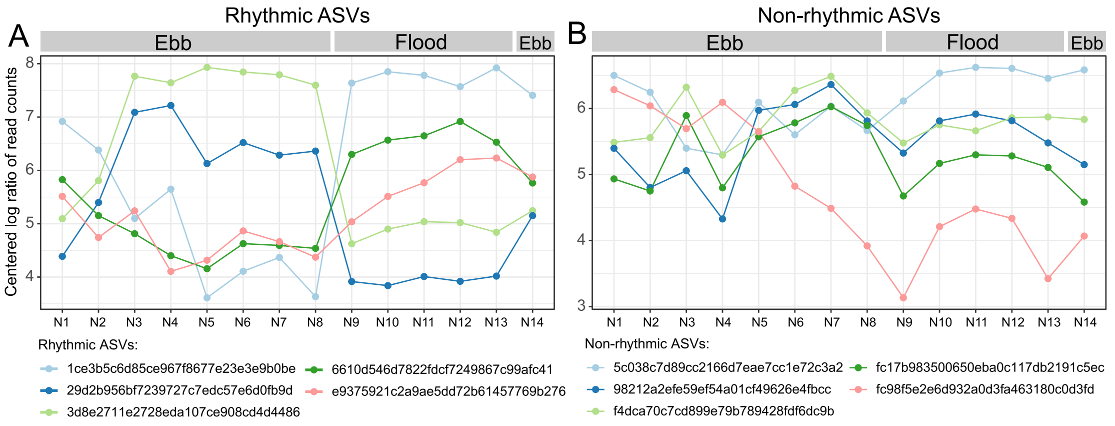


Figure: Temporal abundance (Centered Logarithm Ratio of read counts) trends of representative rhythmic (A) and non-rhythmic (B) ASVs in NTS-Neap. Only top 5 most abundant rhythmic ASVs and non-rhythmic ASVs in NTS-Neap were shown in the figure.

**Text S5: Using rhythmic prokaryotic subcommunities to predict physicochemical variables**

Random forest analysis is an example of machine learning where an ensemble of decision trees are generated to iteratively identify the optimal set of explanatory variables to predict variation in a response variable [8]. Recently. random forest models based on prokaryotic communities have been successfully used to predict physicochemical variables and soil quality [9], similar outcomes have also been reported from the assessment of aquatic communities [10]. Therefore, random forest analyses were performed using the R library 'random forest' with default parameters to predict the physicochemical variables in tidal creek surface water. First, relative abundance table of 399 rhythmic ASVs within 21 (70%) surface water samples was used as the training dataset to construct eight random forest regression models for each physicochemical variable (water depth, salinity, temperature, pH, DO, DIC, DOC and $^{222}Rn$). And these eight models were used to predict their corresponding physicochemical factors based on the training dataset. Second, remaining 8 surface water samples (30%) were used as test dataset to validate the models for each factor. Finally, linear regression analyses between predicated values and observed values for each variable in training dataset and test dataset were used to assess the accuracy of the random forest predictions: $R^2$ and slope values closer to 1 indicated better models (Fig. S8).

**Text S6: Standardized Checkerboard Index analysis for intertidal water microbiota**

The null model analysis of co-occurrence patterns was carried out by Standardized Checkerboard Index analysis (SES C-score) to estimate the species interactions at community level [11-13]. The C-score counts the number of 2 × 2 species matrices where each species occurs only once but at different sites. The standardized effect size (SES) was also obtained to avoiding the influence of various numbers of species and co-occurrences in each crust category on the C-score value. The C-score tests were performed using the 'C-score.R' and 'C-score-var.R' scripts of package MbioAssy1.0 (https://github.com/emblab-westlake/MbioAssy1.0) with the sequential swap randomization algorithm and 30 000 simulations. A positive SES value of C-score indicates segregation between species, and a negative value, aggregation. Specially, C-score value would not differ significantly from that expected at random in the case of similar degree of positive and negative interactions. Therefore, the SES values of $C_{var}$-score were further calculated to assess whether there was a simultaneous increase in both positive and negative interactions, especially for these groups with lower value of SES C-score [14]. In this study, both lowest positive values of SES C-score and SES $C_{var}$-score observed in NTG group indicated weakened degree of species segregation, rather than increased degree of species aggregation (i.e., positive interactions).

**Text S7: Co-occurrence network analysis**

ASVs in at least half of the samples for each network (390 ASVs for HTG, 368 for NTG, 467 for NTS-Ebb, 430 for NTS-Flood, 525 for NTS-Neap, and 395 for NTS-Spring) were included for the correlation calculation. The network-level topological features, including total number of nodes, total number of edges, total number of modules, network density, average clustering coefficient (avgCC), average path distance (GD) and modularity (MD) of each network were calculated by the online Molecular Ecological Network Analysis Pipeline (MENAP) analysis (http://ieg4.rccc.ou.edu/mena/main.cgi).

For each node, its within-module connectivity (Zi) and among-module connectivity (Pi) were calculated in the MENAP and used to determine its topological roles. The keystone nodes including module hubs ($Zi \geqslant 2.5$, $Pi < 0.62$), connectors ($Zi < 2.5$, $Pi \geqslant 0.62$), and network hubs ($Zi \geqslant 2.5$, $Pi \geqslant 0.62$) were identified according to criteria from previous studies [15-17]. All other nodes were categorized as peripheral nodes.

For each network, 100 corresponding Erdös-Réyni random networks were generated with the same network size and average number of links using the 'Random_network.R' script of package MbioAssy1.0 (https://github.com/emblab-westlake/MbioAssy1.0). In addition, the small-world properties of each network were calculated with a coefficient $\sigma = (avgCC/avgCCr)/(GD/GDr)$ and $\sigma > 1$ indicates 'small world' properties, i.e., high interconnectivity and high efficiency. (avgCCr: CCr for random network and GDr: GD for network).

### Supplementary Figures

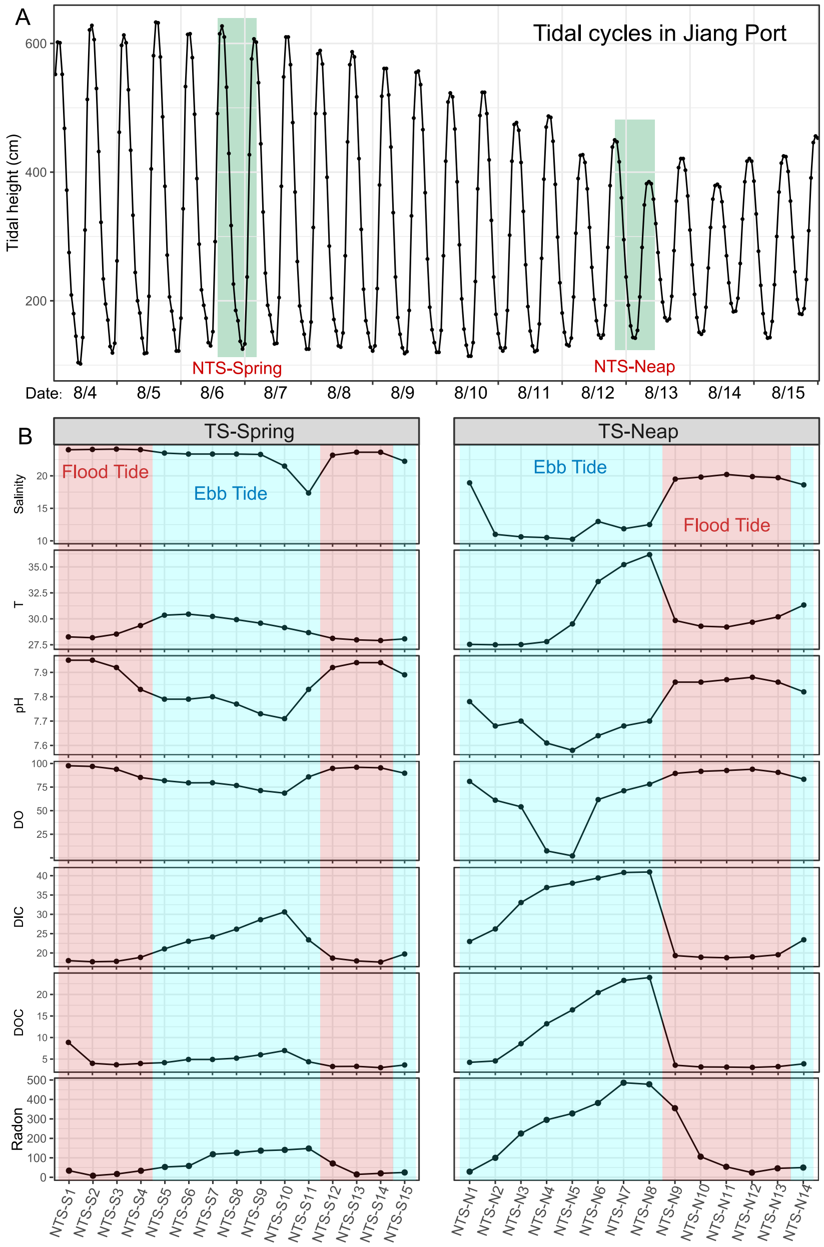


Figure S1. Time series sampling of surface water in the times series station of natural tidal creek (NTS) and temporal dynamics of physicochemical factors. (A) Tidal height of tidal cycles observed in the Jiang Port, where is about 30 kilometers away from the sampling tidal creek. The time periods for collecting surface water at neap tide (14 sample for 13 hours) and spring tide (15 sample for 14 hours) are marked with green rectangle, respectively. (B) Geochemical observations of time-series surface water measured at neap tide and spring tide, including salinity, temperature (T, ℃), pH, dissolved oxygen (DO, %), dissolved inorganic carbon (DIC, mg/L), dissolved organic carbon (DOC, mg/L) and Radon (Bq/m$^3$).

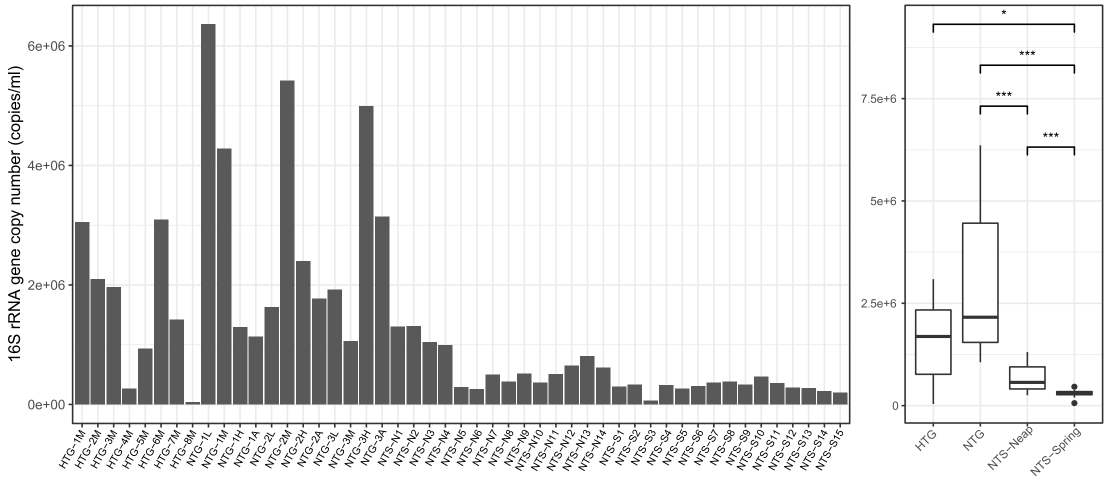


Figure S2. Prokaryotic biomass concentration of groundwater and surface water in Saltmarsh Tidal Creek of Yancheng National Nature Reserve, China. The concentration values were estimated based on 16S rRNA gene copy numbers determined by quantitative PCR and compared across individual water samples (left panel) or their groups (right panel). HTG and NTG represented groundwater (as a proxied by porewater) samples collected from a human-impacted or natural tidal creek, respectively. NTS-Neap and NTS-Spring represented surface water samples collected from time-series station during a neap or spring tide. The significance level in the median differences across groups was statistically checked using Wilcoxon rank-sum test. Only significant differences were shown in right panel Significance was expressed as ****p* < 0.001, ***p* < 0.01, **p* < 0.05.

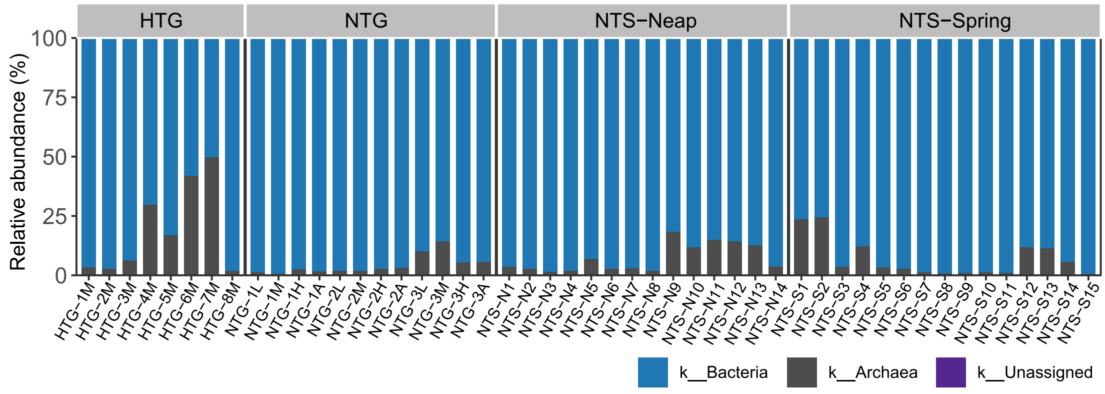


Figure S3. Kingdom-level microbial composition in tidal creek groundwater and surface water. The result showed that archaeal communities were significantly more abundant in the human-influenced tidal creek groundwater (HTG; 1.76%-49.66% and median is 11.56%) than the natural one (NTG; 0.56%-14.25% and median is 2.56%), while their relative abundance all increased from upstream to downstream groundwater. Detailed data in Supplementary Table S4.

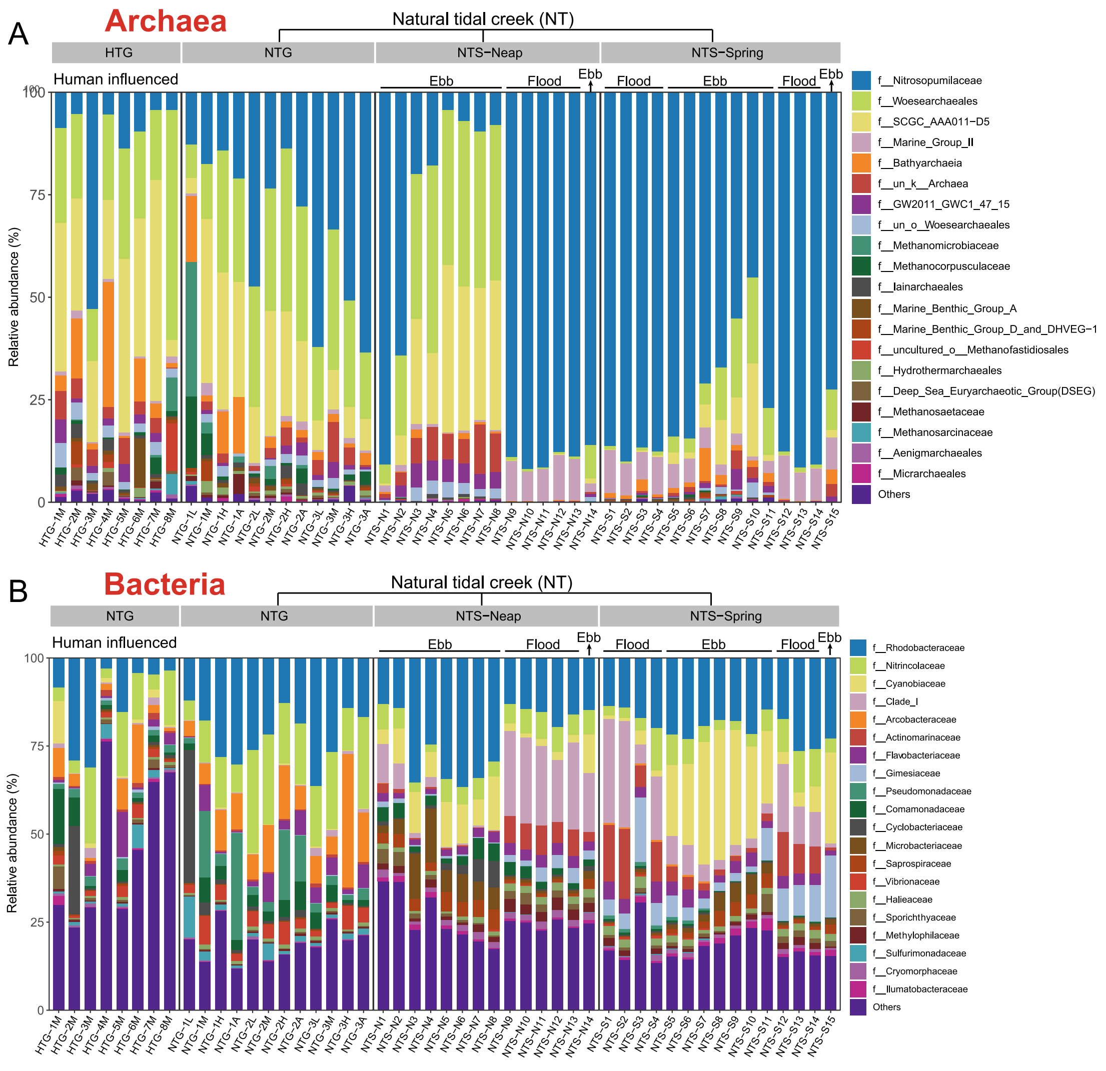


Figure S4. The relative abundance of family-level archaeal (A) and bacterial (B) communities in intertidal groundwater and surface water samples.

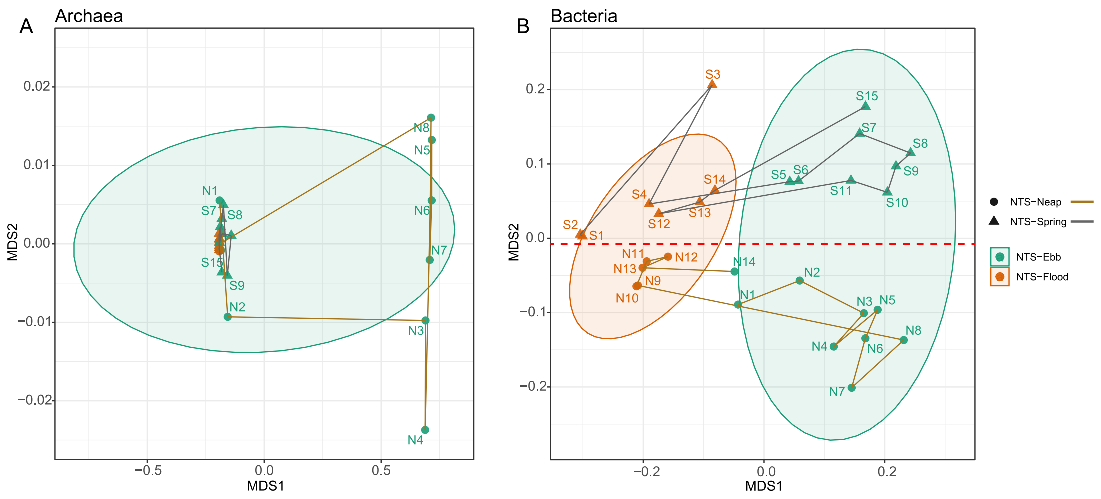


Figure S5. The NMDS ordination of archaeal (A) and bacterial (B) communities in intertidal surface water based on weighted UniFrac distance. The sample trajectories indicated sampling order of surface water during neap tide (N1-N14, brown lines) and spring tide (S1-S15, gray lines). The results showed that the intriguing microbiota circadian rhythm patterns were superimposed by highly synchronized and complex bacterial community dynamics and contrastingly weak (but abrupt) and simpler archaeal community dynamics.

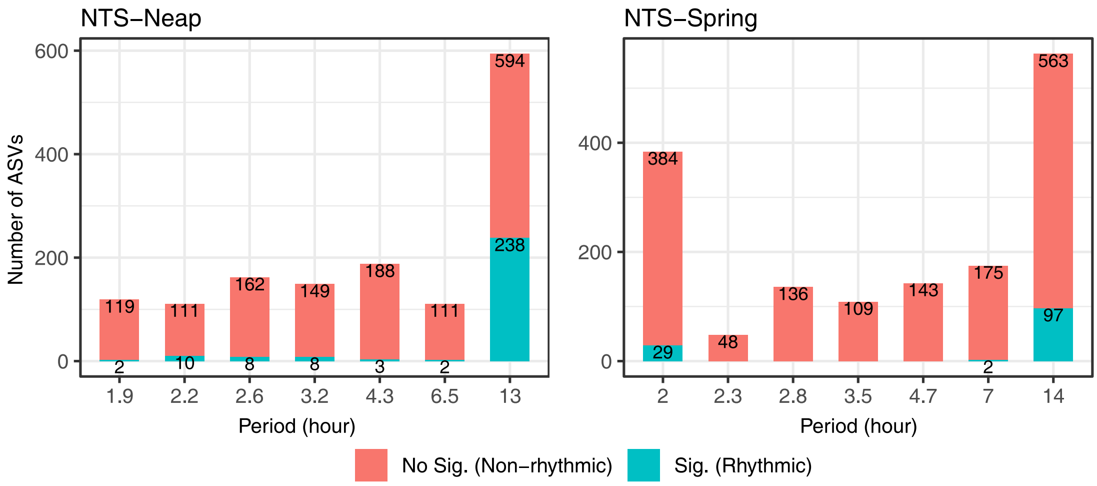


Figure S6. Number of rhythmic ASVs and non-rhythmic ASVs with different rhythmic periods identified in NTS-Neap and NTS-Spring. The number of rhythmic ASVs (statistically significant in LSP analysis with p-value < 0.05) and non-rhythmic ASVs (no significant in LSP analysis with p-value >= 0.05) for each rhythmic period was shown in the graph.

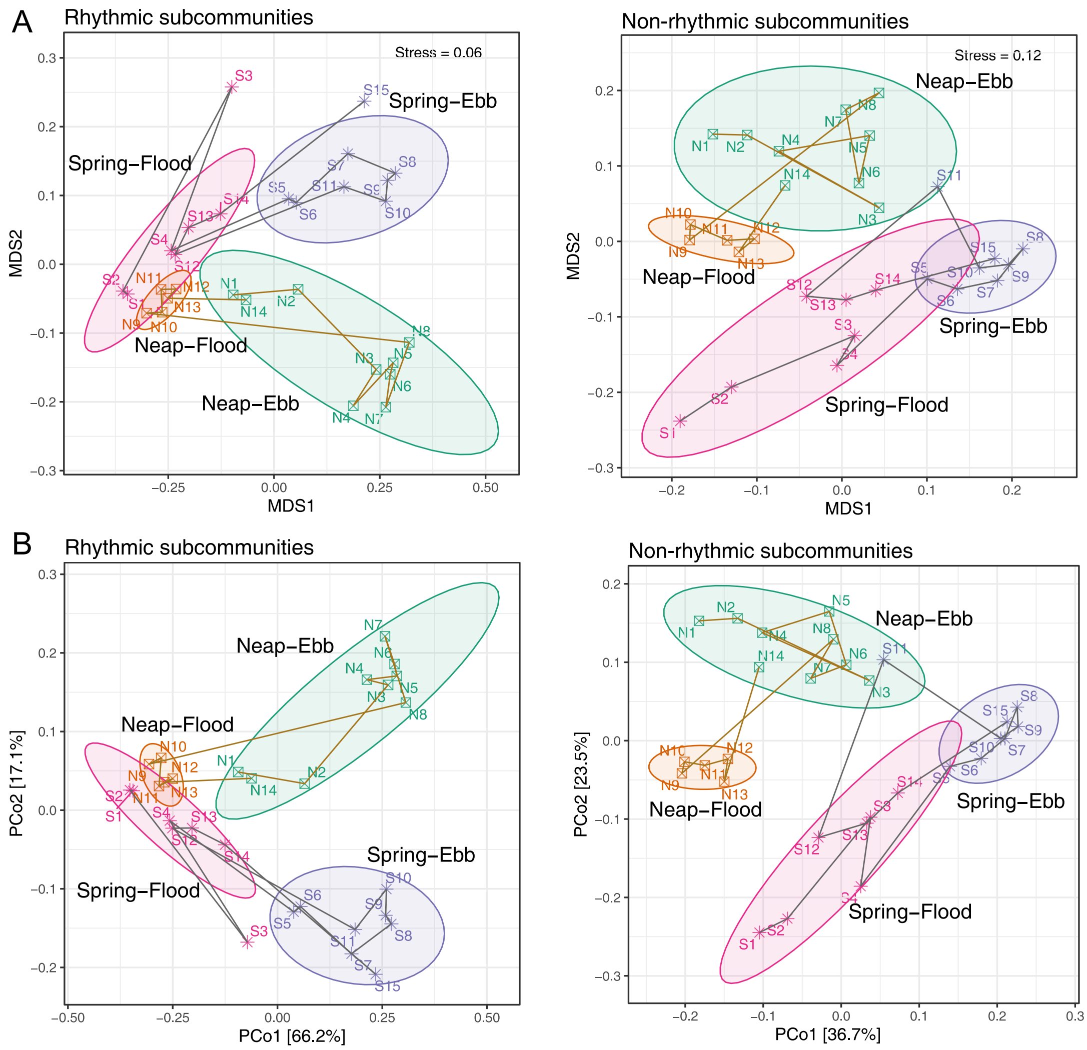


Figure S7 The NMDS (A) and PCoA (B) ordination of rhythmic subcommunities and non-rhythmic subcommunities in intertidal surface water based on weighted UniFrac distance. Rhythmic ASVs identified in NTS-Neap and NTS-Spring via LSP analysis were grouped into rhythmic subcommunities and the other ASVs were grouped into non-rhythmic subcommunities.

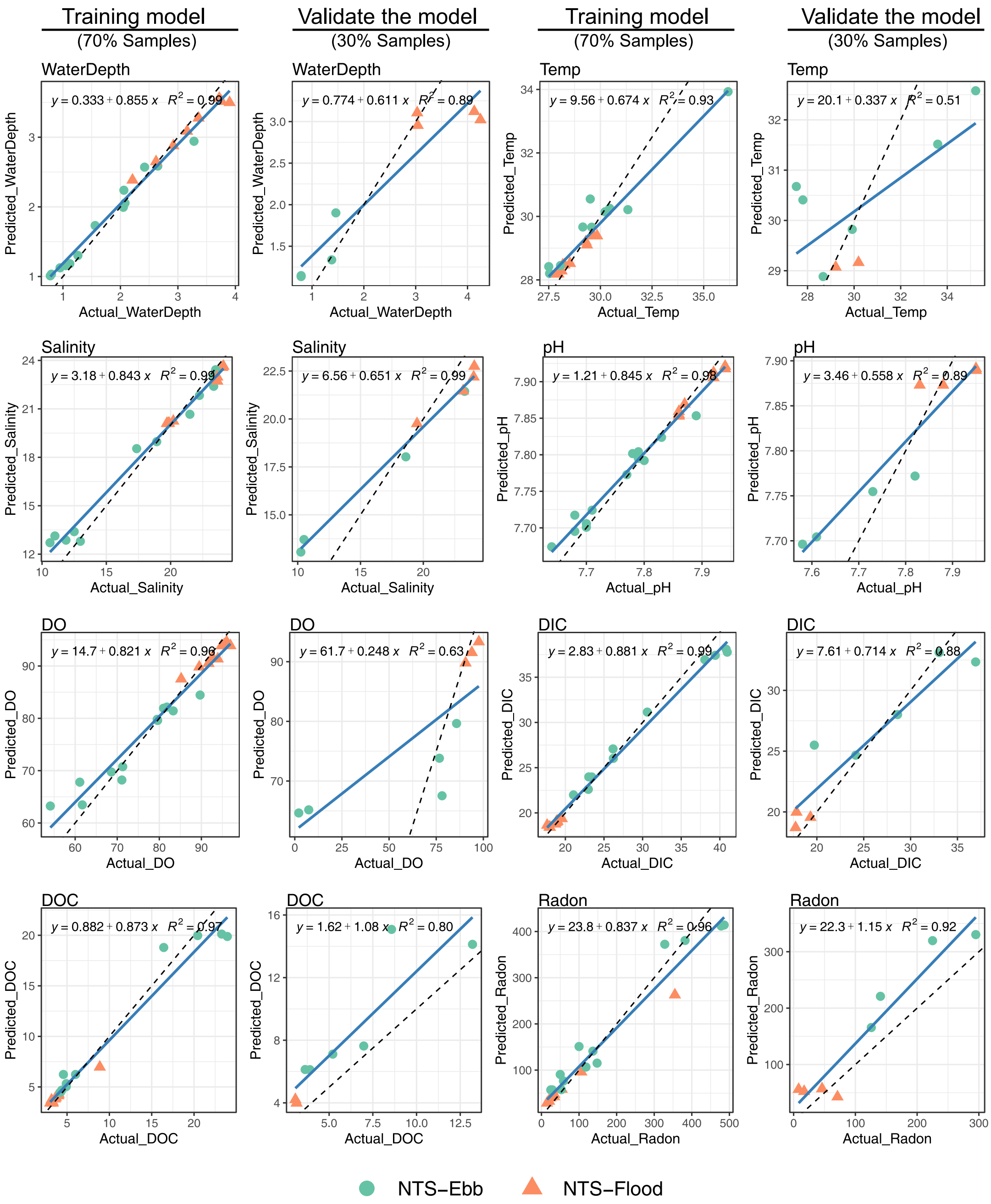


Figure S8. Predicted physicochemical values based on random forest regression model versus actual values. Linear regression analyses between predicated values (random forest models) and observed values of each variable were used to assess the accuracy of the random forest predictions in the training dataset (21 surface water samples, 70%) and test dataset (8 surface water samples, 30%) in NTS group. Dashed black lines indicate where points should fall for a perfect prediction. Blue lines indicate linear fitting of predicted values. Adjusted $R^2$ and fitted model for each linear regression are indicated on the plots and $R^2$ and slope values closer to 1 indicated better models.

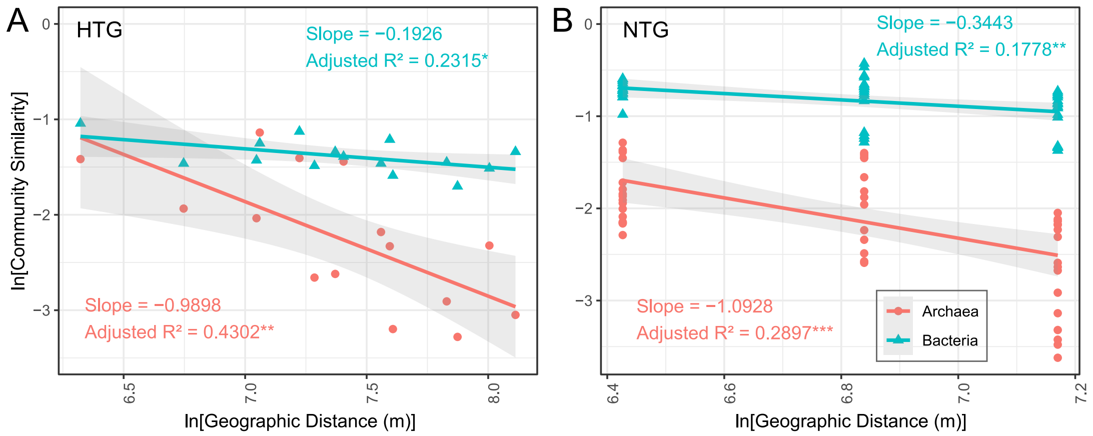


Figure S9. Distance-decay relationships (DDRs) of archaeal (red lines) and bacterial (blue lines) communities. DDRs showing Bray-Curtis similarity against geographic distances between sampling sites in the human influenced (HTG, A) and natural (NTG, B) tide creek, respectively. Solid lines denote the ordinary least-squares linear regressions. The DDRs values are presented as the slope. The $R^2$ were calculated reflecting variance explained by the LM model. Significance was expressed as ****p* < 0.001, ***p* < 0.01, **p* < 0.05.

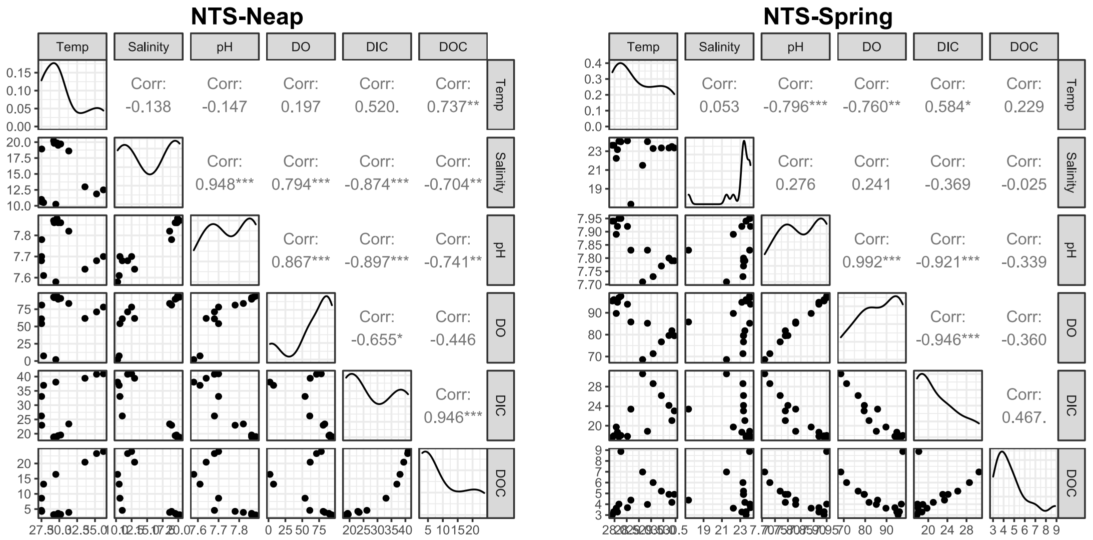


Figure S10. Correlations between each pair of environmental factors observed in intertidal surface water collecting at neap tide and spring tide, including salinity, temperature (Temp, °C), pH, dissolved oxygen (DO, %), dissolved inorganic carbon (DIC, mg $L^{-1}$), and dissolved organic carbon (DOC, mg $L^{-1}$).

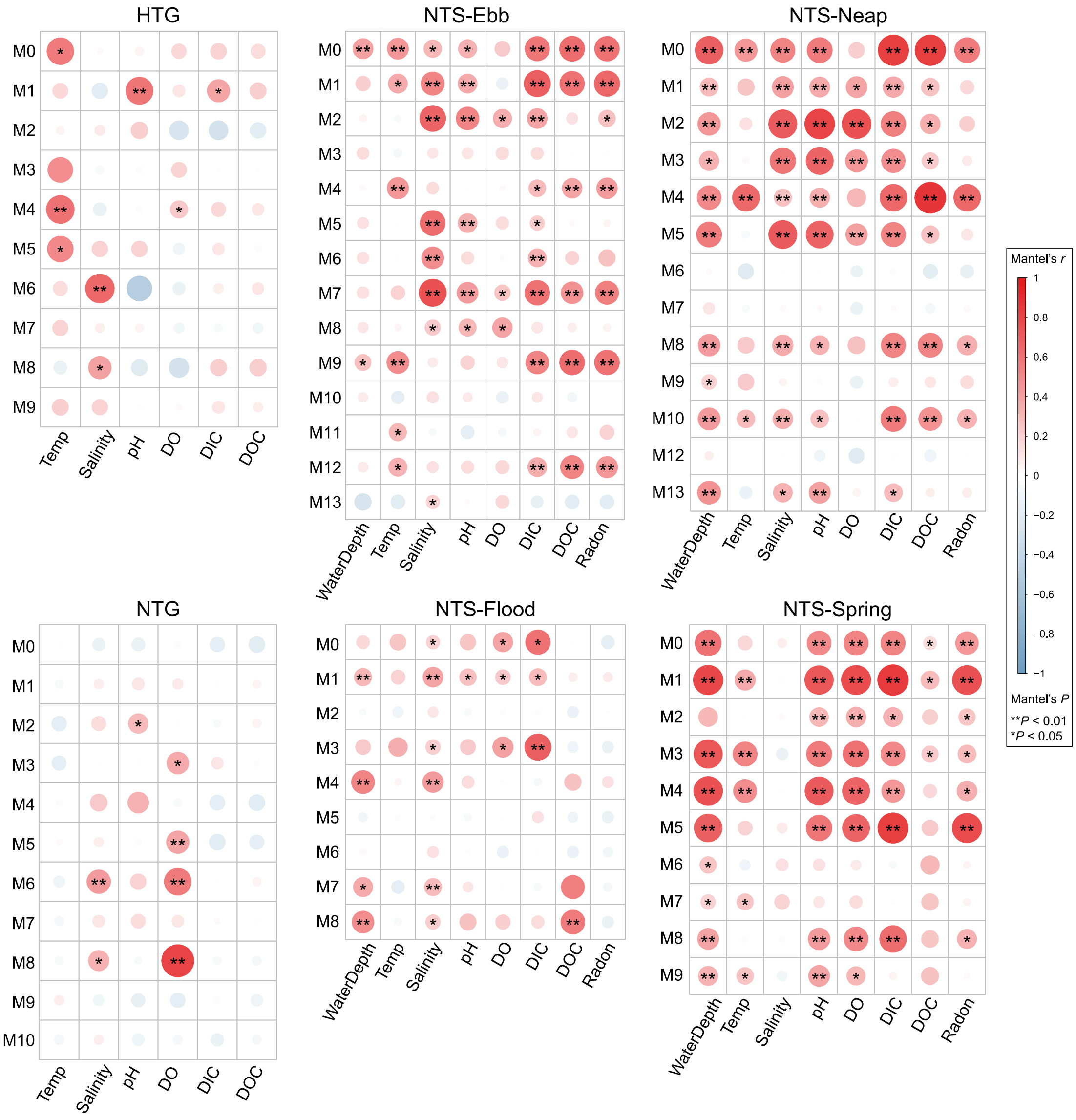


Figure S11. Heatmap plots of correlations between environmental factors and microbiota in each module of each co-occurrence network based on partial mantel tests. Only modules containing at least three ASVs are included. Significance was expressed as $**P < 0.01$, $*P < 0.05$

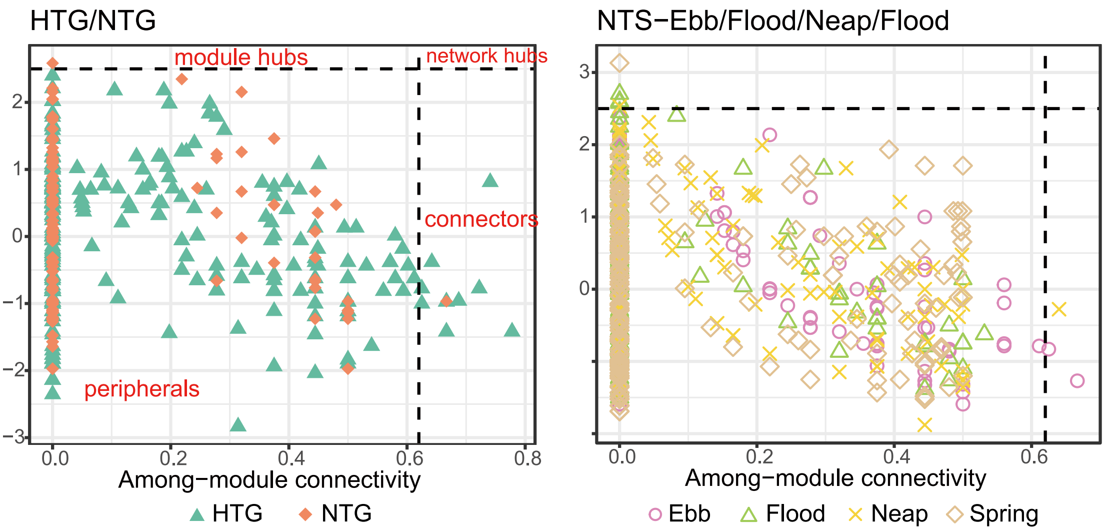


Figure S12. Putative keystone taxa identified based on the node topology in each network. Each symbol represents a node in one of the networks. A node is identified as a module hub if its within-module connectivity (Zi) ⩾ 2.5 and among-module connectivity (Pi) < 0.62, as a connector if its Pi ⩾ 0.62 and Zi < 2.5, as a network hub if its Zi ⩾ 2.5 and Pi ⩾ 0.62, and all other nodes are identified as peripherals (Zi < 2.5 and Pi < 0.62). Detailed information for these keystone taxa is listed in Table S16.

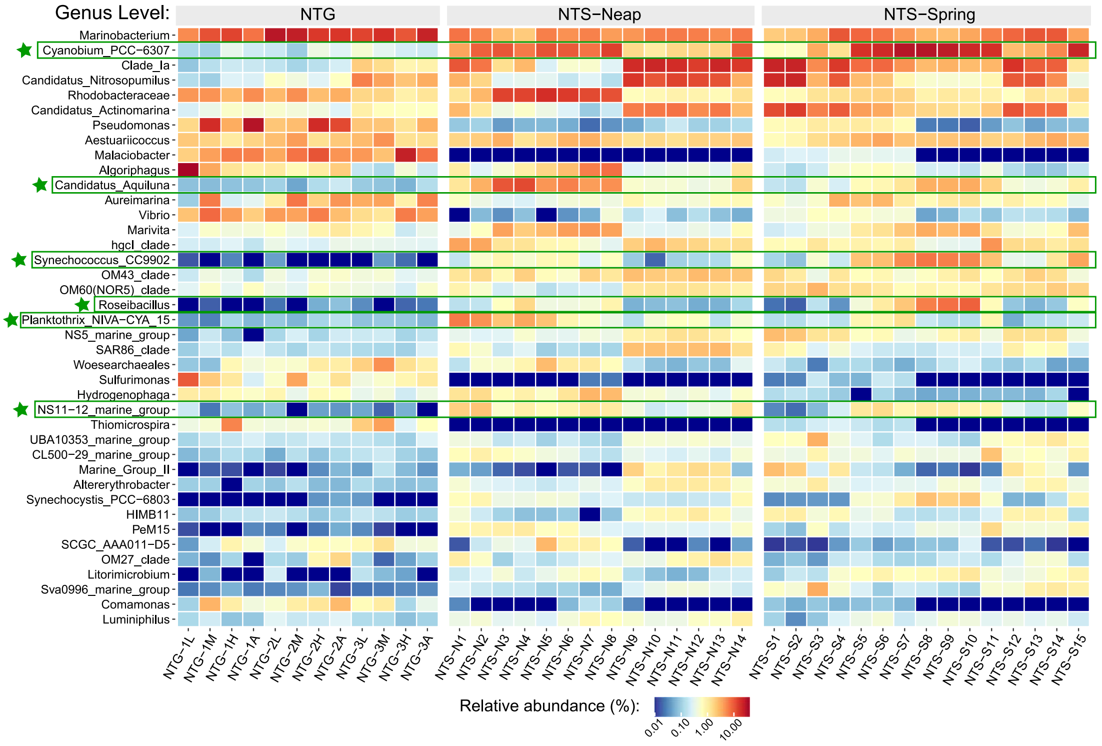


Figure S13. Heatmap plots of relative abundance of genus-level prokaryotic communities in natural intertidal groundwater-surface water continuum. Only top 40 genera with higher relative abundance were showed in this figure. These genera marked in the figure showed higher relative abundance in the ebb-tide surface water than in the intertidal groundwater and flood-tide surface water.

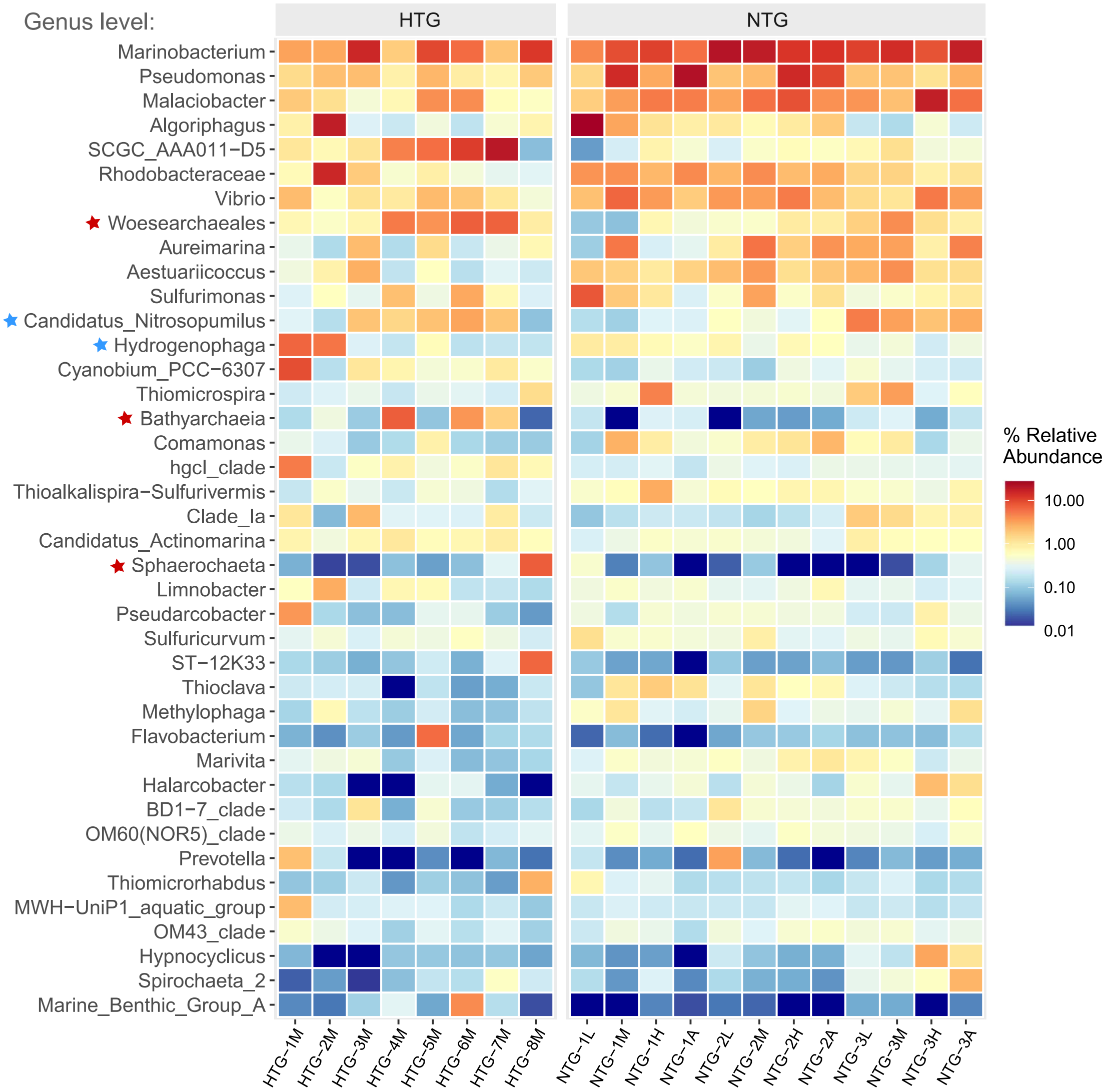


Figure S14. Heatmap plots of relative abundance of genus-level prokaryotic communities in intertidal groundwater. Only top 40 genus with higher relative abundance were showed in this figure. These genera marked in the figure showed higher relative abundance in the human-influenced creek groundwater than in the natural creek groundwater.
